# In-Depth Characterization of Stem Cell Potency and Genotoxicity for Clinical-Scale Ex Vivo CRISPR/Cas9 Gene Editing

**DOI:** 10.64898/2026.01.08.698401

**Authors:** A Naseem, W Vetharoy, T E Whittaker, F Zinghirino, R Ghosh, E Gomez Castaneda, H Ali, C. Cipriani, E. Montini, A J Thrasher, G Santilli, D Cesana, G Turchiano, A Cavazza

## Abstract

Translating CRISPR/Cas9-based homology-directed repair (HDR) strategies into clinical application remains a major challenge due to limited standardization, concerns over safety, and efficacy issues. Here, we present a comprehensive and clinically compliant preclinical framework for the ex vivo correction of Wiskott-Aldrich Syndrome (WAS) using a CRISPR/Cas9-AAV6 platform targeting hematopoietic stem and progenitor cells (HSPCs). In this study, we established a clinical-compatible platform enabling large-scale manufacturing while preserving HSPC viability, stemness, and multilineage functionality. To overcome low HSPC long-term engraftment, we fine-tuned AAV dosing and transiently modulated p53 signalling, achieving significantly improved in vivo correction and repopulation. Importantly, we implemented a multi-tiered genotoxicity assessment strategy, integrating in silico, genome-wide, and orthogonal assays, revealing a largely favorable safety profile with minimal off-target risks and no signs of clonal dominance or transformation. Longitudinal in vivo safety monitoring revealed donor specific rare off-target events and structural variants. This highlights the crucial importance of patient monitoring after transplantation, further emphasized by the identification of a de novo chromosomal rearrangements that could be detected exclusively following cell engraftment in mice. This work offers a robust and adaptable roadmap for future HDR-based gene editing platforms, establishing critical benchmarks for efficacy, safety, and regulatory readiness in the development of advanced therapeutic medicinal products.

## Introduction

The discovery of DNA endonuclease enzymes, especially those linked to the clustered regularly interspaced short palindromic repeats (CRISPR) system, has revolutionized genetic correction in hematopoietic stem and progenitor cells (HSPCs). CRISPR/Cas9 targets specific genomic sites using a guide RNA that pairs with DNA. Once it identifies a sequence adjacent to a protospacer adjacent motif (PAM), it introduces a double-strand break (DSB), which can be repaired by the cell’s natural DNA repair mechanisms, such as non-homologous end joining (NHEJ) or microhomology-mediated end joining. Alternatively, if a therapeutic donor sequence is provided alongside CRISPR machinery, the cell can use homology-directed repair (HDR) to incorporate the corrective sequence (Allen et al. 2023; Bahal et al. 2024; Castiello et al. 2024; Dever et al. 2016; Fananas-Baquero et al. 2021; Rai et al. 2020; Rai et al. 2024). Because many monogenic disorders arise from loss-of-function mutations affecting entire gene loci, insertion of a complete therapeutic coding sequence via HDR offers a universal therapeutic solution while ensuring controlled gene copy number and transcriptional regulation. In this regard, promising preclinical results have been obtained by delivering CRISPR components to HSPCs via electroporation of ribonucleoprotein (RNP) complexes, combined with HDR donor templates delivered through recombinant adeno-associated virus 6 (AAV6) vectors. (Allen et al. 2023; Bahal et al. 2024; Castiello et al. 2024; Dever et al. 2016; Fananas-Baquero et al. 2021; Rai et al. 2020; Rai et al. 2024).

Despite its potential, HDR-based gene editing still faces significant challenges in achieving durable corrections for blood-related genetic diseases and thus in reaching clinical translation. One major hurdle is that current techniques primarily edit committed progenitors rather than long-term repopulating stem cells while also imposing detrimental effects on cell fitness, reducing their viability, frequency and regenerative capacity. Many preclinical studies have reported a decline in hematopoietic repopulation capacity, both in vitro and following in vivo transplantation. This reduction is primarily attributed to the activation of a p53-mediated DNA damage response and innate immune signalling triggered by Cas9 activity and donor template delivery, particularly when using AAV vectors. These cellular responses drive hematopoietic stem cell differentiation and contribute to the exhaustion of their long-term repopulating potential (Della Volpe et al. 2024; Ferrari et al. 2022; Lee et al. 2024; Schiroli et al. 2019). As such, efforts to improve both the frequency of corrected primitive hematopoietic stem cells (HSCs) and their engraftment in mice upon in vivo transplantation have led to significant results when using optimized protocols to transiently manipulate DNA repair pathways, cell cycle and culture conditions to favour HDR or to dampen detrimental signaling pathways (Baik et al. 2024; Conti et al. 2025; De Ravin et al. 2021; Rai et al. 2023). On the other hand, unintended genetic modifications, RNA editing, integration of portions of donor templates, and chromosomal abnormalities at both on- and off-target sites are potential side effects that still pose a major concern for clinical application of gene editing. Numerous technologies have emerged to assess the safety profiles of gene editors using both qualitative and quantitative assays in vitro and in living cells (Blattner et al. 2020). However, the field has yet to establish standardized protocols to evaluate the genotoxic risks associated with different gene-editing techniques. Additionally, the lack of reliable methods to predict the functional consequences of genomic alterations makes it difficult to interpret the vast amount of safety data generated with respect to their clinical relevance. Therefore, there is an urgent need to establish a consensus on the key aspects that should be addressed in preclinical studies of HSPC gene editing platforms to enable their clinical testing and eventual commercialization (Jeffers et al. 2024).

Previously, we showed that HDR-mediated insertion of a correct copy of the *WAS* gene, involved in the occurrence of a rare primary immunodeficiency – Wiskott-Aldrich Syndrome (WAS) – via CRISPR/Cas9 and AAV6 in patient-derived HSPCs led to full correction of all functional hallmarks of the disease *in vitro* and *in vivo* (Rai et al. 2020). The platform reached curative levels of gene knock-in, leading to restored physiological expression of the *WAS* gene, functional correction of all hematopoietic lineages, absence of altered HSPC functionality and controlled gene integration at the *WAS* locus with normal gene copy number. Here, we present a series of preclinical experiments that define and refine our genome editing strategy, bringing it closer to clinical trial readiness. This set of experiments comprehensively investigate the adoption of a GMP-grade scale up protocol for editing HSPCs while assessing the functionality of manipulated stem cells and the toxicity and safety of the whole procedure by longitudinal analysis *in vitro* and *in vivo*. Overall, our preclinical studies suggest that HDR-mediated editing of HSPCs at the *WAS* locus is feasible and reproducible at clinical scale, with sustained cell viability and repopulating capacity when optimized protocols are implemented. Thorough safety analysis confirmed absence of genotoxic events caused by the gene editing procedure and highlighted the necessity of patient-tailored assessments and post transplantation evaluations of functional relevance of potential off target editing. Moreover, this study lays the foundation for developing a clear roadmap of essential preclinical tests required to support the clinical translation of HDR-based gene editing platforms in HSPCs for the treatment of a broad spectrum of blood disorders.

## Results

### Optimization of HSPC gene editing protocols to cGMP-compliant standards

To set up a cGMP-compliant HSPC manipulation pipeline, we fine-tuned our previously developed gene editing protocols to match the standards required for a clinical-grade manufacturing process. First, we modified the AAV6 donor vector carrying a codon optimized WAS cDNA flanked by *WAS* homology arms (Rai et al. 2020) by switching from Ampicillin to Kanamycin resistance in the vector backbone and by shortening the homology arms from ∼750 bp to ∼400 bp for improved AAV production and detection of the knock-in cassette in target cells and tissues (**Figure S1A**). By comparing both donor vector designs in *in vitro* knock-in experiments in HSPCs, we confirmed consistent HDR efficiency when using the new construct. Then, we replaced research-grade cell culture and gene editing reagents with GMP-grade ones, again showing stable HDR rates when using the new set of reagents (**Figure S1B**). Optimization of Cas9:gRNA stoichiometry to reduce overall manufacturing costs as well as to calibrate gRNA complexation with Cas9, resulted in 1:1.65 being the most cost-effective and efficient Cas9:gRNA ratio in our setting, in terms of knock-in efficiency and cell viability (**Figure S1C**). Lastly, we tested the use of the cGMP-compatible MaxCyte GTx electroporator by comparing the use of different electroporation programs to previously used parameters in the Neon NTx electroporation system. By employing the program HSC3 in the MaxCyte GTx machine, we achieved highly consistent HDR rates in HSPCs, paralleled by superior cell viability (**Figure S1D, E**).

Having established a reliable GMP-grade platform to edit HSPCs, we then moved forward to scale up HSPC manipulation to clinical need using the pipeline shown in **Figure 1A**. CD34+ HSPCs were isolated from 8 batches of G-CSF mobilized peripheral blood from healthy donors either manually or using the automated CliniMACS Prodigy system, with successful and reproducible cell purification (**Figure 1B and Figure S2**). Once purified, HSPCs were pre-stimulated for 2 days in a defined medium supplemented with cytokines, followed by Cas9:gRNA ribonucleoprotein (RNP) electroporation and transduction with the AAV6 coWAS donor. A portion of edited cells, alongside controls, was then harvested at day 4 for quality control (QC) testing, such as viability, editing efficiency (Indel/HDR) as well as in vitro genotoxicity analysis. In parallel, the majority of cells were instead cryopreserved, followed by thawing, QC analysis and transplantation into animals, to mimic a potential clinical process.

**Figure 1.**
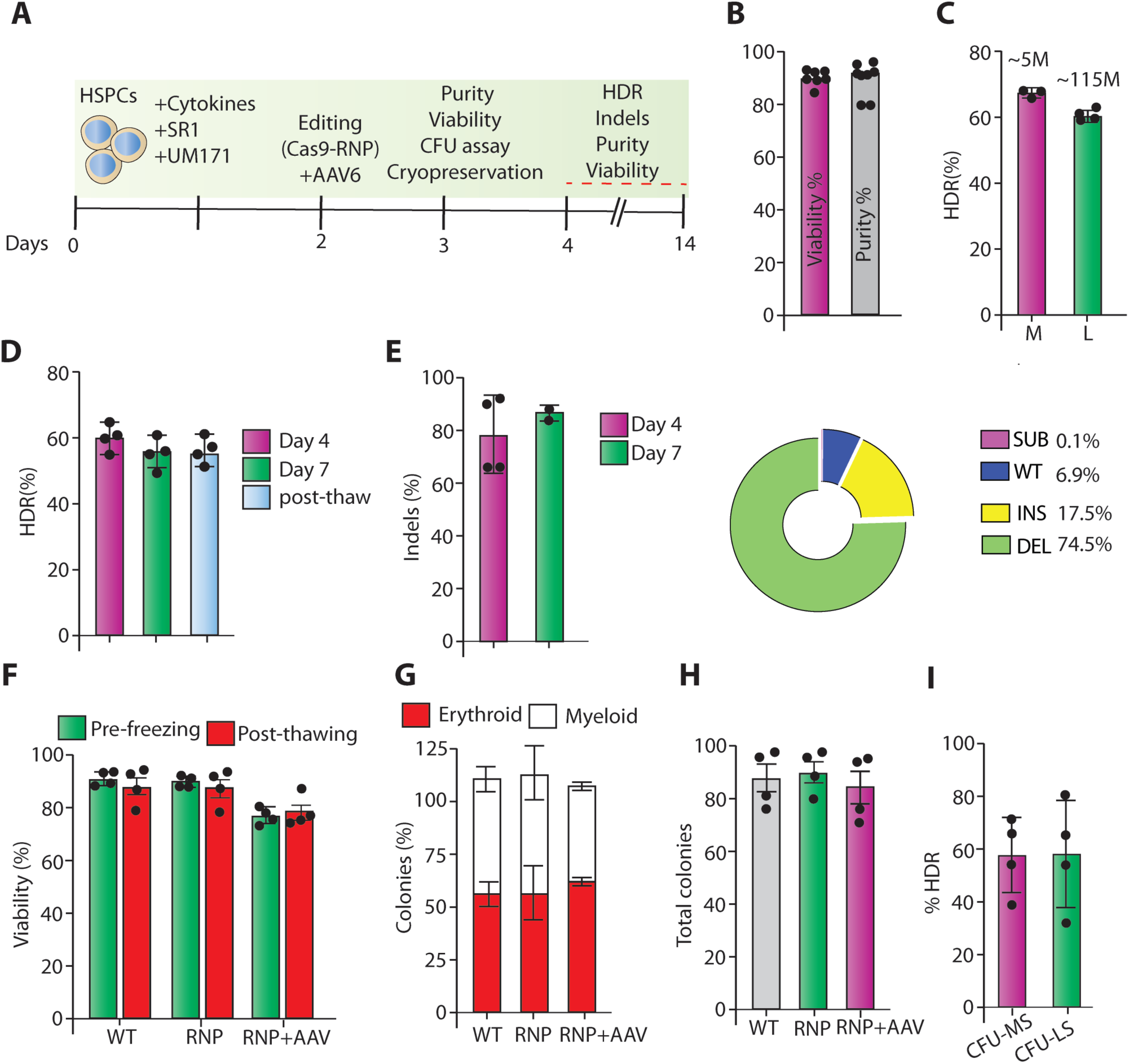
Feasibility of large-scale HSPC gene editing under cGMP-compliant conditions. **(A)** Schematic overview of the experimental workflow and timeline. **(B)** Viability and purity (CD34⁺ cell content) of hematopoietic stem and progenitor cells (HSPCs) isolated from leukapheresis derived from healthy donors (n=7 biological replicates), assessed by flow cytometry. **(C)** Knock-in efficiency of the WAS cDNA cassette in HSPCs from medium-scale (n = 4 biological replicates) and large-scale (n = 4 biological replicates) experiments, determined by droplet digital PCR (ddPCR). **(D)** Knock-in efficiency in large-scale edited HSPCs as assessed by ddPCR at day 4 and day 7 post editing and after cryopreservation. **(E)** Bar graph showing insertion/deletion (InDel) frequencies in genomic DNA from RNP-edited HSPCs at day 4 and day 14 post-editing, determined by next-generation sequencing (NGS). The pie chart on the right displays the distribution of predominant mutations identified by NGS. **(F)** Viability of manipulated (RNP and RNP+AAV) and unmanipulated (WT) HSPCs, evaluated before and after cryopreservation, as assessed via DAPI staining and flow cytometry (n = 4 biological replicates). **(G)** Bar graphs showing the percentage of myeloid (white) and erythroid (red) colonies formed in semi-solid cultures by wild-type (WT), RNP-edited, and RNP+AAV-edited/transduced HSPCs from healthy donors (n = 4 biological replicates). **(H)** Total number of colonies generated in solid cultures as described in (G). **(I)** Frequency of WAS cassette knock-in assessed in colonies generated in semisolid cultures by RNP+AAV HSPCs edited at large (LS) and medium (MS) scale (n=4 biological replicates). Data are presented as mean ± SD.

Targeted integration analysis by droplet digital PCR (ddPCR) confirmed insertion of the coWAS cassette in 61% of CD34⁺ cells when editing an average of 115 × 10⁶ HSPCs, and in 67% of cells when using a medium-scale setup involving 5 × 10⁶ cells per experimental replicate. (**Figure 1C**). HDR rates were stable throughout the *in vitro* culture up to 14 days after editing and in the cryopreserved product upon defrosting and infusion into animals (**Figure 1D**). Indel rates in electroporated samples (RNP) reached up to 95% (range 67-95%) and again remained stable throughout the in vitro culture (**Figure 1E**). This underscores absence of toxicity or negative selection of large scale manipulated cells in vitro, further confirmed by >75% of cell viability detected both before and after cryopreservation of the edited cell product, with negligible differences compared to unmanipulated or RNP controls (**Figure 1F**; **Figure S3**). Importantly, unmanipulated (WT) and manipulated (RNP and RNP+AAV) HSPCs showed comparable clonogenic potential in solid culture, with no significant differences in the overall quantity of Colony Forming Units (CFU) and no evidence of lineage skewing among the experimental groups (**Figure 1G, H**). Assessment of HDR frequency in randomly picked single CFU colonies showed an average rate of 59% of coWAS targeted integration in HSPCs edited at both large and medium scale (**Figure 1I**), suggesting efficient editing of cells with repopulating capacity. Overall, these results show that our developed GMP-compliant protocol for clinical scale manufacturing of gene edited HSPCs met all the required specifications for cell product potency (>50% gene correction), cell purity (>70% CD34+), cell viability (>70%) and colony forming capacity in solid cultures (>50 colonies every 500 plated cells).

### Robust engraftment of HSPCs manufactured at large scale

To test the ability of the manufactured product to engraft long term and reconstitute haematopoiesis, a total of 250 million large-scale edited and cryopreserved HSPCs from 4 different healthy donors (SU-1 to SU-4) were injected intravenously into sub-lethally irradiated non-obese diabetic–scid-gamma (NSG) mice, alongside WT and RNP controls (as detailed in **Figure 2A&B** and **Material and Methods**). Tail vein bleed 8 weeks post infusion showed comparable human chimerism across the 3 groups (**Figure S4A**). Evaluation of long-term engraftment potential and hematopoietic reconstitution, together with HDR and indel frequencies within the CD45+ human cell compartment, was carried out 12-14 weeks after transplantation. Overall, ex-vivo cultured and/or edited HSPCs showed robust engraftment capacity, with a median human chimerism in the BM of around 39% (range 3-68%), 35% (range 8-61%) and 30% (range 4-65%) for WT, RNP and RNP-AAV, respectively (**Figure 2C**). The human graft in the BM was multilineage and consisted of mostly CD19+ B cells and CD33+ myeloid cells, as usually seen in NSG mice (**Figure 2D and Figure S4B**). We also observed comparable chimerism in the human CD34+ HSPC, multipotent progenitor (MPP) and HSC compartments among experimental groups (**Figure 2E** and **Figure S5A**). While the percentage of engraftment in the BM for the large-scale manufactured products was slightly, although non-significantly, decreased compared to unmanipulated cells, we observed comparable engraftment rates in the periphery (**Figure 2F-G**). DdPCR analysis on sorted human CD45+ cells from the BM of transplanted mice revealed an average of 39.4% and 10.1% of Indels and HDR-mediated coWAS knock-in, respectively (**Figure 2H**), with a similar trend being also observed in CD45+ cells collected from spleen, peripheral blood (PB) and thymus (**Figure S5B**), indicating engraftment of edited HSPCs and their correct differentiation in all the hematopoietic organs.

**Figure 2.**
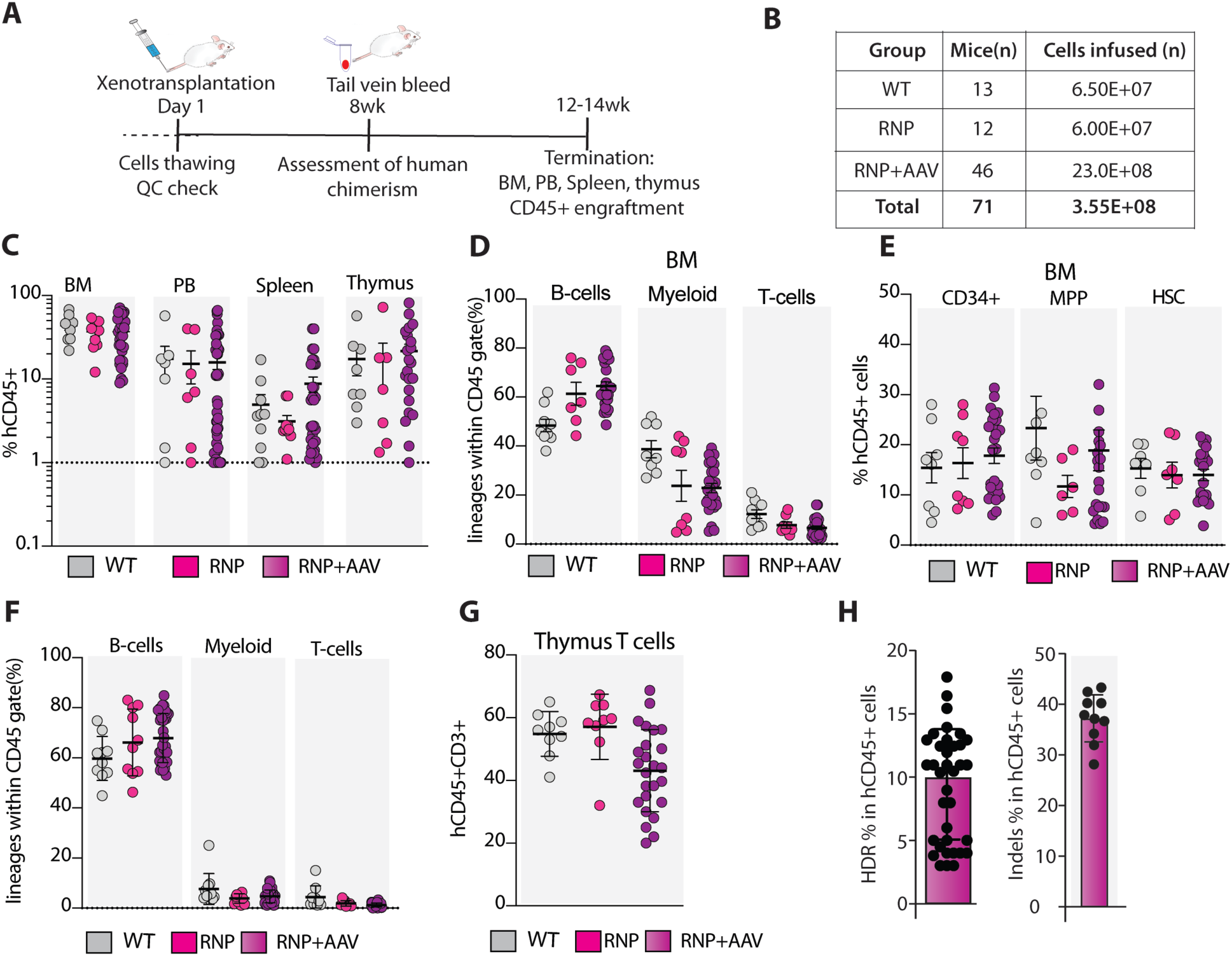
Effective engraftment and multilineage differentiation of large-scale gene-edited HSPCs in vivo. **(A)** Schematic representation of the in vivo experimental design using the NSG mouse model. **(B)** Summary table indicating the total number of HSPCs injected and the number of recipient animals per experimental group. **(C)** Human chimerism levels, expressed as percentages of CD45+ cells, in bone marrow (BM), spleen, peripheral blood (PB), and thymus of transplanted NSG mice, assessed 12 weeks post-transplantation. Each dot represent data retrieved from one mouse. **(D)** Proportional distribution of human hematopoietic lineages (B cells, T cells, and myeloid cells) within the human (hCD45⁺) cell compartment in the BM. Each dot represent data retrieved from one mouse. **(E)** Distribution of human stem and progenitor subsets—multipotent progenitors (MPP), hematopoietic stem cells (HSCs), and CD34⁺ cells—within the BM human cell population in transplanted mice. Each dot represent data retrieved from one mouse. **(F)** Lineage distribution of human hematopoietic cells in the spleen of transplanted mice, as described in (D). Each dot represent data retrieved from one mouse. **(G)** Frequency of CD3⁺ T cells within the hCD45⁺ population in the thymi of transplanted mice across all experimental groups. Each dot represent data retrieved from one mouse. **(H)** Quantification of WAS cassette integration (HDR) and InDel formation in human CD45+ cells isolated from the BM of mice transplanted with RNP+AAV HSPCs, measured by ddPCR and Sanger sequencing, respectively. Each dot represent data retrieved from one mouse. Data are presented as mean ± SD, except for panels C-E were mean ± SD was used.

The decrease in the engraftment frequency of HSPCs corrected via HDR over the course of long-term transplantation studies in mice and in patients is a well-known phenomenon, largely ascribed to poor stem cell editing and procedure-related cell toxicity. We therefore sought to investigate the impact of our large-scale gene editing protocol on HSPC gene expression profile by comparing the transcriptome of unmanipulated, electroporated-only (RNP) and fully gene edited (RNP+AAV) HSPCs collected 2 days after editing, the usual time point for product infusion into patients when treated via gene therapy. Principal Component Analysis (PCA) highlighted distinct expression profiles imputable mainly to the different donor sources (PC1 and PC2), suggesting minimal influence on the overall transcriptome by the gene editing procedure; only 16% of the variance was indeed attributed to cell manipulation (PC3) and only in the presence of AAV transduction (**Figure S6A**). This is reflected by the low number of differentially expressed genes (DEGs) retrieved, with 149 and 26 DEGs in AAV+RNP when compared to WT and RNP, respectively (FDR<0.05; FC>2). No differential gene expression was observed when comparing WT versus RNP condition, underscoring the lack of toxicity caused by the electroporation procedure optimized in this study (**Figure 3B and Figure S6B, C**). Since all DEGs emerging from the AAV+RNP *versus* RNP comparison were shared with the AAV+RNP *versus* WT comparison (**Figure S6D**), we focussed on the latter and performed a functional enrichment analysis to investigate the presence of pathways and biological processes related to the transcriptome change after HDR-mediated gene editing (**Figure 3A**). Genes differentially expressed in the RNP+AAV condition belonged to very interconnected categories in the MSigDB database, including cell cycle, apoptosis, cell stress, interleukin signalling, DNA repair and TP53 signalling (**Figure 3B**). Interestingly, 14/72 genes upregulated in the RNP+AAV condition were direct transcriptional targets of TP53, underscoring the presence of a stress-related HSPC response to the use of AAV donors for HDR, as previously observed (Ferrari et al. 2020; Schiroli et al. 2019).

**Figure 3.**
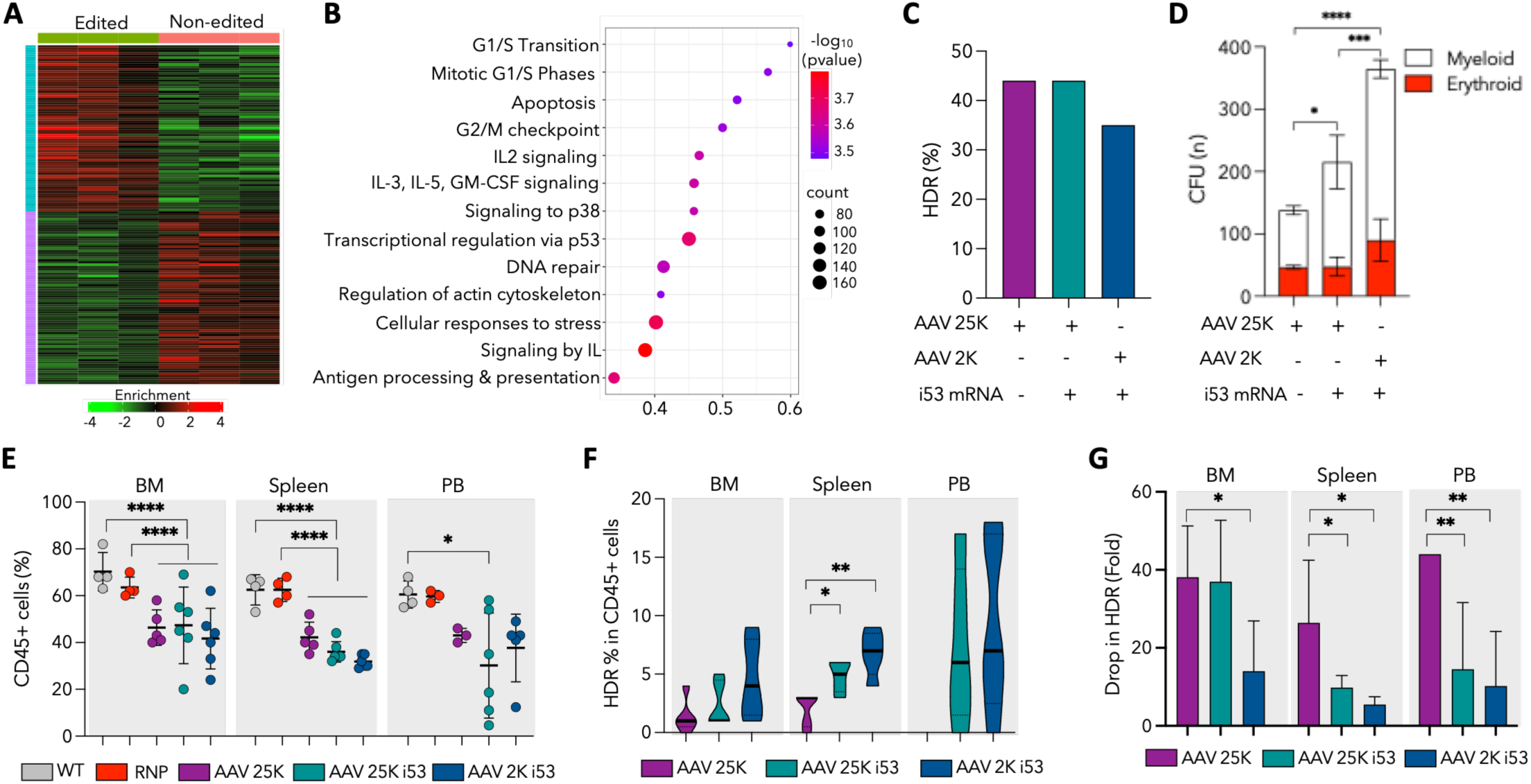
Dampening of p53 signalling improves engraftment of HSPCs that underwent HDR. **(A)** Representative hierarchical clustering map of differentially expressed genes when comparing unedited (WT) and edited (RNP+AAV) HSPCs transcriptomes (n=3 biological replicates). The blue bar indicates the cluster of genes upregulated in edited HSPCs, while the purple bar indicates the cluster of genes downregulated in edited HSPCs. **(B)** Pathway enrichment bubble plot, representing the pathways most enriched in differentially expressed genes when comparing unedited (WT) and edited (RNP+AAV) HSPCs transcriptomes. P values are represented by bubble color, gene counts are represented by bubble size. **(C)** Frequency of HDR observed in RNP+AAV HSPCs edited using different AAV MOIs with or without the addition of i53 mRNA. **(D)** Bar graphs showing the percentage of myeloid (white) and erythroid (red) colonies formed in semi solid cultures by RNP+AAV HSPCs edited using different conditions as described in (C) (\*\*\*\**p* < 0.0001, *** *p* < 0.001, \**p* < 0.05; 2-way ANOVA with Tukey multiple comparison test) **(E)** Human chimerism levels, expressed as percentages of CD45+ cells, in bone marrow (BM), spleen and peripheral blood (PB) of transplanted NSG mice, assessed 12 weeks post-transplantation. Each dot represent data retrieved from one mouse (n=4-6 mice per experimental group; \*\*\*\**p* < 0.0001, \**p* < 0.05; 1-way ANOVA with Tukey multiple comparison test). **(F)** Quantification of WAS cassette integration (HDR) in human CD45+ cells isolated from the BM, spleen and PB of mice transplanted with RNP+AAV HSPCs edited using different conditions, measured by ddPCR. Each dot represent data retrieved from one mouse (n=4-6 mice per experimental group; \*\**p* < 0.01, \**p* < 0.05; 2-way ANOVA with Tukey multiple comparison test). **(G)** Bar plot representing the drop in HDR frequency post-transplantation for each experimental condition analysed. The drop was calculated as fold decrease in HDR frequency observed for each experimental mice (n=4-6 mice per experimental group; \*\**p* < 0.01, \**p* < 0.05; 2-way ANOVA with Tukey multiple comparison test) at 12 weeks post-transplantation in CD45+ cells in the BM, spleen and PB, compared to pre-transplant HDR rates observed in HSPCs for each experimental condition (C). Data are presented as mean ± SD.

In light of these evidences, and with the aim to increase the frequency of engraftment of HSPCs with a correct coWAS knock-in in the BM of recipient mice, we modified the manufacturing protocol to include the use of an inhibitor of 53BP1, which has been shown to dampen TP53-mediated signalling and thus gene editing-mediated cell death and toxicity, (Baik et al. 2024; De Ravin et al. 2021) and optimized AAV doses via a Response Surface Methodology approach, (Whittaker et al. 2024) to achieve efficient knock-in frequencies while preserving cell fitness. HSPCs from different healthy donors were electroporated with RNPs and transduced with the AAV donor vector under three conditions: at a multiplicity of infection (MOI) of 25,000 with or without i53 mRNA (referred to as AAV 25K and AAV 25K i53, respectively), and at a MOI of 2,000 with i53 mRNA (AAV 2K i53). All manipulations were performed within the framework of our optimized large-scale manufacturing protocol. Culture and manipulation of HSPCs with the presence of i53 mRNA greatly reduced p21 expression irrespective of the AAV doses used (**Figure 6E** and **Figure S6E**). DdPCR revealed correct knock-in of the coWAS cassette in the *WAS* locus, with an HDR rate of 45% in AAV 25K and AAV25K i53, which slightly decreased to 35% when scaling down AAV dose by 10 fold (**Figure 3C**). Interestingly, the addition of i53 to the culture medium resulted in a similar frequency of HDR events, without any further enhancement in gene knock-in efficiency, in contrast to previous reports (Baik et al. 2024; De Ravin et al. 2021). HSPCs placed in solid cultures gave rise to a significantly increased numbers of colonies when i53 was supplemented to the culture media, and the effect was further and significantly enhanced when combined with reduced AAV doses, with almost a 4-fold increase in colony numbers compared to the AAV 25k condition (**Figure 3D**). Two days post editing, cells were transplanted into NSG mice alongside unmanipulated controls and engraftment was assessed 12 weeks later. We observed an average of 45% (range 42-47%) engraftment in the BM and 37% (range 32-43%) in peripheral hematopoietic organs in all experimental conditions where AAV was employed, with a small but significant decrease compared to unmanipulated and RNP controls, especially in the BM and in the spleen (**Figure 3E**). DdPCR analysis on human CD45+ cells sorted from the BM, PB and spleen of transplanted mice showed a visible trend towards increased engraftment of RNP+AAV manipulated cells when i53 mRNA was included in the culture media, which was even more pronounced when combined with lower AAV doses (**Figure 3F**). If compared to pre-transplant HDR rates, AAV 2K i53 HSPCs displayed the lowest drop in HDR post-transplant in all hematopoietic organs analysed (average of 7.6, 5 and 3.8 fold decrease in the BM, spleen and PB respectively) followed by AAV 25K i53 (average of 18, 9.1 and 5.9 fold decrease in the BM, spleen and PB respectively) and AAV 25K (average of 31.4, 22 and 44 fold decrease in in the BM, spleen and PB respectively) (**Figure 3G**), further confirming the detrimental effect that AAV donors exert on edited HSPCs and the benefit of fine tuning multiple manipulation conditions to achieve better therapeutic outcomes.

### No Detectable In Vivo Toxicity After Transplantation of HSPCs Produced at Clinical Scale

To assess the long-term potential for in vivo toxicity and tumorigenesis as a result of transplantation of large-scale edited HSPCs, we closely monitored transplanted mice for the development of adverse clinical signs over the 12 weeks following HSPCs infusion (**Figure 4A**). From a toxicology point of view, no mice experienced weight loss after bone marrow conditioning and HSPCs transplantation and overall, all mice gained weight regularly and survived until the end of the experiment (**Figure 4B**). Peripheral blood was collected at termination for haematological analysis, showing no significant difference in the complete blood count and leukocyte differential among experimental groups, suggesting absence of major toxic effects (**Figure 4C, D**). Histopathological analysis of BM smears and spleen sections of 55 transplanted mice carried out in certified GLP conditions confirmed presence of all precursor/progenitor lineages as well as absence of any gross lesions or adverse effects attributable to cellular infusion. Furthermore, no evidence of morphologically disrupted haematopoiesis was reported in all the mice samples analysed, confirming absence of tumorigenesis (**Figure 4E, F and Figure S7A**).

**Figure 4.**
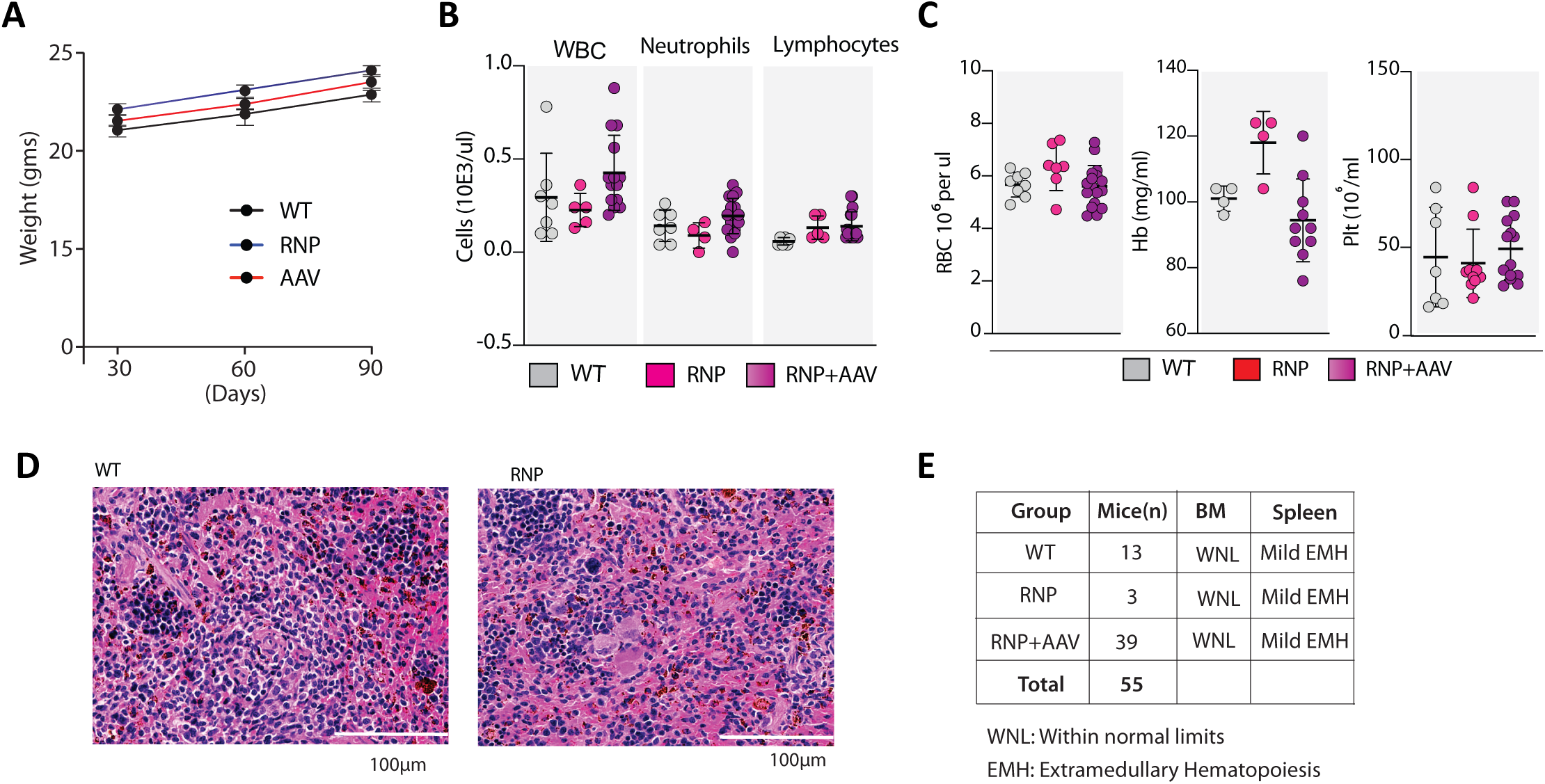
No evidence of in vivo toxicity upon transplantation of HSPCs manufactured at clinical scale. **(A)** Body weight measurements of NSG mice from transplantation through to the experimental endpoint. **(C–D)** Hematological parameters (white blood cells, red blood cells, and platelets) measured in peripheral blood of xenotransplanted NSG mice at 12 weeks post-injection. Each dot represent data retrieved from one mouse. **(E)** Representative images from histopathological analysis of spleen sections stained with hematoxylin and eosin (H&E), captured at 20× magnification. Images shown were taken from mice transplanted with unmanipulated (WT) or edited (RNP+AAV) HSPCs. **(F)** Summary table of histopathological evaluations from bone marrow (BM) smears and spleen sections. BM smears demonstrated expected cellular morphology and normal levels of hematopoietic precursors across all groups. Mild to minimal extramedullary hematopoiesis (EMH) observed in spleen sections was consistent across experimental groups. Data are presented as mean ± SD.

### Extensive Genotoxicity Assessment Demonstrates Safety of the Gene Editing Platform

A key objective of this preclinical study is to assess the safety of the gene editing platform by ensuring efficient on-target genetic correction while minimizing off-target effects. Indeed, one of the primary risks associated with CRISPR technology is the potential for unintended genetic modifications and rearrangements at both on- and off-target sites, which could introduce mutations that may lead to oncogenic transformation—posing a significant concern for clinical applications involving engineered nucleases. (Lee, Lozano, and Dunbar 2021). In this study, we have taken a comprehensive approach for evaluating the safety and toxicity of our gene edited product manufactured at scale, using a combination of DNA and RNA sequencing-based analyses and pathology assays in a time course fashion, both *in vitro* and *in vivo* (**Figure 5A**). Sampling and off-target analysis of samples collected at different time points and especially before and after transplantation allows not only the identification of undesired genetic modifications in the product, but also the assessment of their functional relevance, such as their impact on selection of cells bearing specific off-target modifications or their potential to develop malignancies, or unbalanced haematopoiesis.

**Figure 5.**
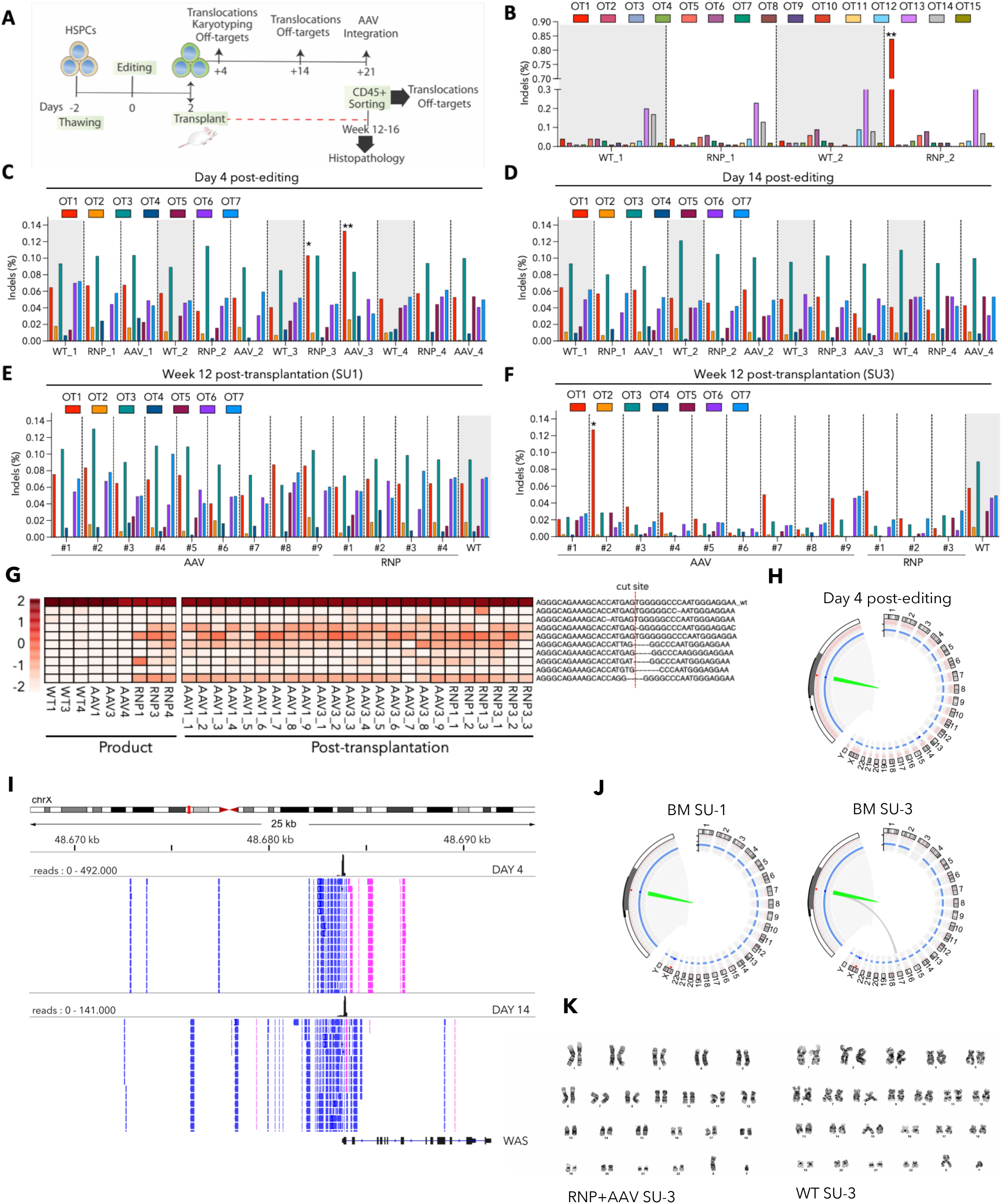
Comprehensive and longitudinal genotoxicity assessment confirms overall safety of the gene editing platform in vitro and in vivo. **(A)** Schematic representation and timeline of the safety study, describing the different assays used to assess genotoxicity in vitro and in vivo. **(B)** InDel frequency analysis in WAS patient-derived HSPCs at 15 potential off-target sites retrieved by in silico prediction (n=2 biological replicates, each utilized in 2 experimental replicates; \**p* < 0.05, Statistical significance was assessed using Z-tests corrected by standard deviation calculated from untreated WT controls). **(C–D)** InDel frequency analysis at on-target and predicted off-target sites, performed by targeted amplicon deep sequencing of genomic DNA (gDNA) isolated from edited (RNP and RNP+AAV) HSPCs at day 4 (**C**) and day 14 (**D**) post-editing (n=4 biological replicates, each utilized in 2 experimental replicates; \*\**p* < 0.01,\**p* < 0.05, statistical significance was assessed using Z-tests corrected by standard deviation calculated from untreated WT controls). **(E–F)** InDel frequency analysis at on-target and predicted off-target sites, performed by targeted amplicon deep sequencing of genomic DNA (gDNA) isolated from CD45+ cells in the BM of mice transplanted with edited (RNP and RNP+AAV) HSPCs from SU-1 (n=14 mice analysed) (**E**) or SU-3 (n=13 mice analysed) (**F**) (\**p* < 0.05, Statistical significance was assessed using Z-tests corrected by standard deviation calculated from untreated WT controls). **(G)** Mutational tracking of gene-edited cells pre- and post-transplantation, showing the relative abundance of the top 10 most frequent alleles at the on-target cleavage site. **(H)** Representative Circos plot of CAST-seq analysis of edited cells at day 4 post-editing, illustrating lack of detection of chromosomal translocations. On-target InDels are shown in green. **(I)** Qualitative CAST-Seq analysis of HSPCs at day 4 and 14 post editing. Representative Integrative Genomics Viewer (IGV) plots illustrate CAST-Seq reads surrounding the target site within a window of 25 kb. Mapped CAST-Seq reads are represented by bars (only the top 7 lines are shown). Blue and pink bars indicate sequences aligning with the negative or positive strand, respectively. Coverage (i.e., the number of mapped reads) is indicated by histogram plots at the top of the bar plots for each experimental condition. Reads mapped to the left side of the cut site indicate large deletions, while those on the right side indicate inversions or/and translocations with homologous regions. **(J)** Circos plots representing CAST-seq analysis of CD45+ cells isolated from the BM of NSG mice transplanted with edited (RNP and RNP+AAV) or WT HSPCs from SU-1 and SU-3. CD45+ cells isolated from from n=2 mice per scale-up experiment (SU1 and SU3) were pooled, each utilized in 2 experimental replicates. In SU-3, the plot highlights on-target InDels (green) and a single translocation to chromosome 17 (gray). **(K)** Representative karyotype of HSPC metaphase spreads analyzed at day 4 post-editing. A total of 270 metaphases from three independent donors were examined, with no chromosomal abnormalities detected. Data are presented as mean ± SD.

To first detect off-target sites in gene edited HSPC products, we performed deep sequencing of 15 off target sites retrieved using the in silico COSMID software based on sequence similarity with the gRNA and previously screened in WAS patient samples (Rai et al. 2020). After validation by NGS, only one site returned significant, although at low frequency, indels in *COL23A1* intron 5 in one HSPC donor source (named as OT1) and as such it was chosen for off-target analysis in large scale edited cells (**Figure 5B**). In parallel, further 6 genomic sites retrieved by an unbiased genome wide GUIDE-seq analysis (Rai et al. 2020) were added to the pool of potential off target loci to be screened in the large-scale products (OT2-OT7) (**Table S1**). Importantly, only one potential off target site was reproducibly retrieved by both platforms. We performed deep sequencing of seven genomic sites on gDNA extracted from large-scale edited CD34⁺ cells (SU-1 to SU-4) four days post-editing. Significant indels were detected only at the OT1 site, and only in one of the four donor-derived scale-up products treated with either RNP alone or RNP+AAV, when compared to unmanipulated WT controls. (**Figure 5C**). When the same sites were analysed on cells cultured for one additional week (day 14 post editing) we could not detect unwanted genetic modification at any of the sites interrogated, suggesting lack of any selection of cells bearing indels at sites other than *WAS* (**Figure 5D**). To evaluate the emergence of new off-target modifications or the persistence of those observed in vitro – indicative of their potential functional relevance – we performed deep sequencing of the seven predicted off-target sites on human CD45⁺ cells isolated from the bone marrow of mice 12 weeks after transplantation with SU-1 to SU-4 HSPCs. Confirming the data obtained at day 14, the latest time point of in vitro culture, indels at off target sites were not retrieved in most of engrafted cells **(Figure 5E**); however, we surprisingly did detect indels at the OT1 site from cells retrieved from the BM of one of the 13 mice transplanted with RNP+AAV CD34+ cells from SU-3, despite lack of detection pre-transplant (**Figure 5F**). In parallel, we also performed a mutational tracking analysis of gene edited cells before and after transplantation by assessing the frequency of the top 10 most abundant allele variants at the on-target gRNA cutting site. While in WT and scaled-up RNP+AAV samples we detected mostly alleles with *WAS* wild type sequence, reflecting absence of manipulation and correct coWAS knock-in respectively, RNP samples exhibited enrichment in specific allele variants resulting from NHEJ-mediated DSB repair, which were consistent among donor replicates (**Figure 5G**). Tracking of these variants in BM samples 12 weeks after transplantation revealed similar pattern of allele frequencies, highlighting absence of selection of clones bearing specific indels at the *WAS* locus while confirming engraftment of HSPCs that underwent on-target gene editing.

Off-target CRISPR/Cas-induced modifications represent just one of many chromosomal rearrangements that can result from targeted gene editing and thus thorough investigation of possible deletions, translocations and gross chromosomal disruptions at on-and off-target sites is mandatory to provide a comprehensive view of the safety of the approach. To assess this, we performed CAST-seq analysis on edited HSPCs at day 4 and day 14 post editing using the methodology described previously (Turchiano et al. 2021). CAST-seq technology can detect on- and off-target translocations down to a frequency of 1 in 10,000 cells and searches for genome wide translocations in an unbiased way. By analysing all manufactured products at both day 4 and day 14 after editing, we could not detect any major chromosomal aberration, further confirming absence of consistent DSB formation at off-target sites with a second, unbiased technology (**Figure 5H**). We could only observe rare events of large deletions and inversion happening at the on-target WAS genomic locus as a consequence of NHEJ-repair; however, as already demonstrated for other sites (Turchiano et al. 2021) and as suggested by our mutational tracking analysis, the frequency of these events significantly declined at later culture time point, especially for the inversions, indicating counterselection of cells bearing these rearrangements (**Figure 5I)**. To get a thorough understanding of the functional relevance of large deletions/ translocations not detected in the product but that might have appeared in vivo upon transplantation, we performed CAST-seq analysis on genomic DNA retrieved from CD45+ engrafted cells in the BM of mice infused with the SU-1 and SU-3 products. While no chromosomal rearrangements were observed in samples retrieved from SU-1-infused mice, we detected a translocation between the on-target site and chromosome 17 in 2 SU-3-infused mice (**Figure 5J**). The site on chromosome 17 mapped to the 13^th^ intron of the solute carrier family 47 member 2 (*SLC47A2*) gene, which is not a known oncogene. Because the site was neither an off-target nor contained putative gRNA sequences or regions significantly homologous (at least 25bp of homology within a 5-kb window) to the on target site, this translocation was considered to be prompted by a naturally-occurring break site, likely caused by the strong DNA replication challenge imposed on HSPCs after their transplantation (Flach et al. 2014; Mehta and Haber 2014). Importantly, deep sequencing confirmed that the translocation happened at very low frequency (**Table S2**) further verifying the safety profile of the platform *in vitro* and most importantly *in vivo*. Karyotype analysis of HSPCs edited at large scale cultured up to 4 days post editing did not reveal any alteration in ploidy or gross chromosomal structural modifications, further confirming the CAST-seq observations (**Figure 5K and Figure S7B**). In summary, these data highlight the overall safety of our genome editing strategy applied to HSPCs, while underscoring the importance of careful, patient-tailored and continuous assessment of the consequences of CRISPR/Cas activity post transplantation.

### AAV donor vector is preferentially trapped at the WAS locus and marks bone marrow repopulating HSPCs

Because our gene editing approach provides HDR-mediated integration of the corrective cassette via AAV transduction, and because many recent reports have documented AAV integration into the host genome and possible related genotoxicity in murine models, we undertook unbiased genome-wide retrieval of AAV6 integration sites (IS) in pre and post transplantation samples using an established methodology and bioinformatic pipeline (Calabria et al. 2023). We analyzed genomic DNA from the 4 *ex vivo* edited scale-up products and from BM samples of 4 mice transplanted with SU-1 and SU-3, which were carefully selected based on high engraftment rate of HDR-corrected cells. AAV-genome junctions were amplified using three sets of primers, to identify integration via different portions of the AAV genome (**Figure S8A**). In the ex-vivo dataset, we identified 115 non-redundant IS, ranging from 18 to 39 unique IS per large scale editing replicate, while in the post transplantation samples a total of 64 IS were retrieved, with only 9 found in SU-3 transplanted mice, possibly reflecting the lower rate of engraftment in those mice compared to SU-1ones (**Figure 6A, Figure S8B, Table S3**). Integrations were found in enriched regions (**Figure 6B; Figure S8C, D**). The preferential target for integration was *WAS,* accounting for 9.8% of all the integrations retrieved in the analysed samples, followed by *SMPDL3B*, which was targeted by 4.3% of integrations (**Figure 6C and Table S4**). The IS in *WAS* were either found clustered around the gRNA targeting site, indicative of AAV trapping at the Cas-induced DSB, or around *WAS* endogenous regions also contained in the AAV donor vector homology arms, suggesting integration via homology between vector and host genomic sequences (**Figure 6D**). These observations were further confirmed when inspecting the regions of AAV found within the amplified AAV-genome junctions; indeed, while all IS mapped close to the gRNA on target site contained intact 3’ and 5’ ITRs, indicative of vector trapping within DSBs, the remaining IS in *WAS* displayed a pattern similar to the IS retrieved genome-wide, and involved AAV regions such as the polyA, vector backbone or the right homology arm (**Figure 6E**). Finally, none of the IS co-mapped with off target sites either detected by GUIDE-seq or predicted *in silico*, further validating the specificity of our Cas9:gRNA platform.

**Figure 6.**
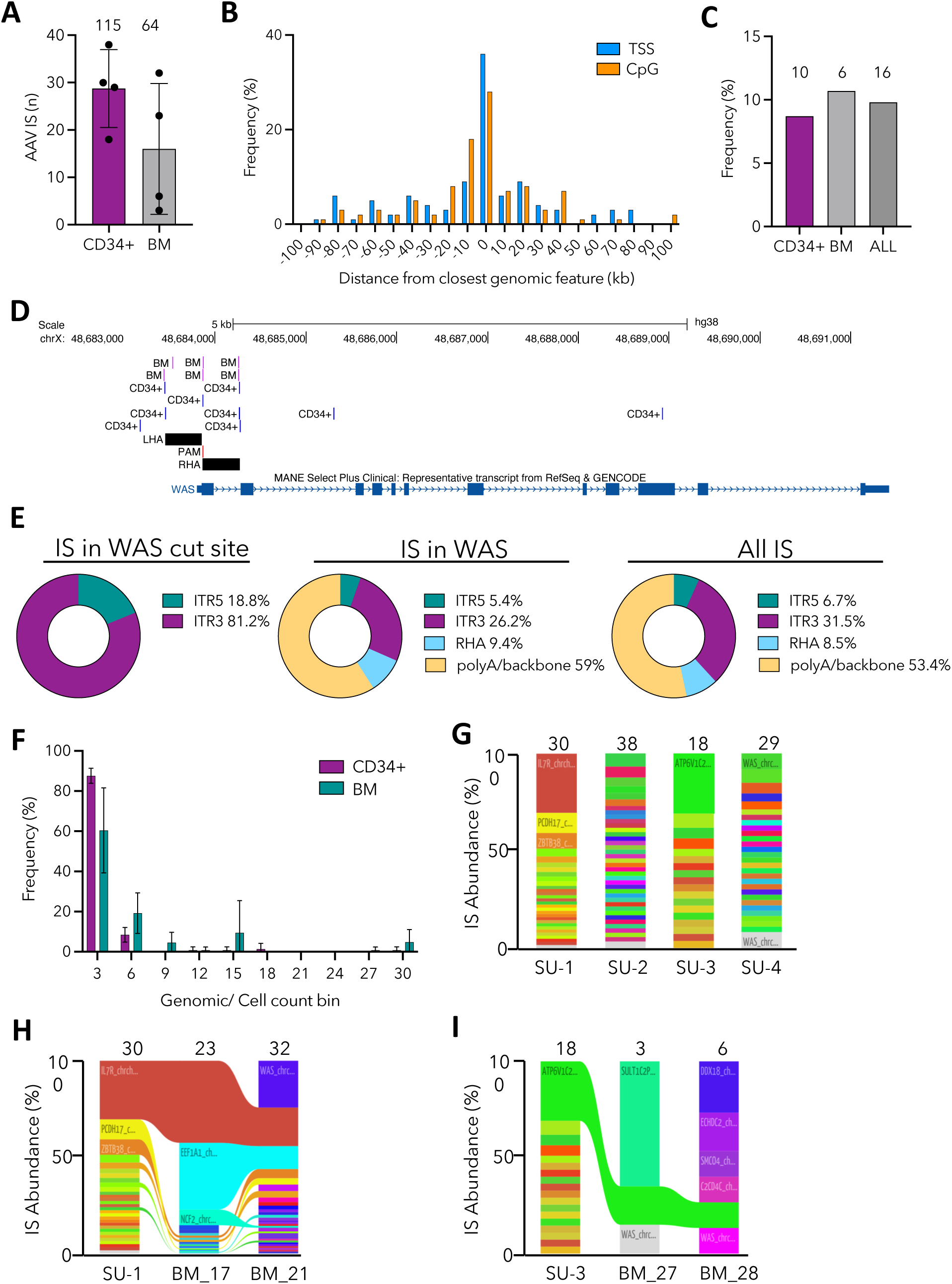
Comprehensive assessment of AAV HDR donor integration in gene edited HSPCs pre- and post-transplantation. **(A)** Number of AAV integration retrieved in the scaled-up products (SU-1 to SU-4) in vitro and in the BM of mice transplanted with SU-1 and SU-3 HSPCs. **(B)** AAV integration site distribution around transcription start sites (TSS) and CpG island. **(C)** Frequency of AAV integrations retrieved within or near the *WAS* locus in in vitro samples (CD34+), in vivo samples (BM) and in all (ALL) the samples analysed. The number of unique IS identified in each category is indicated above each column.. **(D)** Representative UCSC Genome Browser snapshot of the *WAS* locus. AAV integration retrieved from edited HSPCs in vitro (blue bars) and from CD45+ cells isolated from the BM of transplanted mice (pink bars) are depicted. The gRNA PAM site (red bar) and the left and right homology arms contained in the AAV donor molecule (LHA and RHA, black bars) are also indicated. **(E)** Pie charts indicating the frequency of AAV features found at the junction breakpoint between the integrated vector and the host genomic region for the different IS data sets as indicated. **(F)** Bar plot indicating the number of cellular genomes observed for each integration site (Genomes/IS) for each dataset (in vitro CD34+ and in vivo BM) are presented as indicated. **(G-I)** Stacked bar plots showing the relative abundance of AAV IS retrieved in HSPCs edited at large scale (G) and in the BM of mice transplanted with SU-1 (H) and SU-3 (I) HSPCs. In each column, every integration is represented by a different color whose height is proportional to the number of genomes retrieved for that specific IS (percentage of IS abundance, y-axis). The number of unique IS identified in each sample/mouse is indicated above each column. Ribbons connect AAV ISs tracked between samples retrieved before (SU) and after (BM) transplantation. Data are presented as mean ± SD.

The employed AAV integration analysis pipeline makes use of fragmented genomic DNA, enabling the estimation of AAV IS abundance by counting the distinct genomic breakpoints (Calabria et al. 2023). More than 80% and 70% of IS in ex-vivo edited HSPCs and in BM samples respectively were represented by up to 3 genomes/cells, indicating absence of clonal expansion of AAV IS-bearing cells when cultured *in vitro* for up to 21 days post editing or when transplanted in vivo (**Figure 6F**). After estimating the number of genomes linked to each AAV IS, we determined the relative abundance of each IS as a percentage of the total IS identified in each sample. An overall polyclonal pattern was observed in all 4 HSPC scale up samples (**Figure 6G and Table S5**), although clones with a relatively higher level of abundance could be identified in the post transplantation samples because of the small numbers of IS recovered, such as in the case of BM samples from SU-3 transplanted mice (**Figure 6H, I and Table S6**). Nevertheless, tracking of IS abundance before and after transplantation clearly demonstrated engraftment and long-term repopulating potential of gene edited HSPCs, with many IS-bearing clones being shared between ex vivo HSPC and BM retrieved samples (**Figure 6H, I and Table S7**). Analysis of shared IS between pre- and post-transplant samples also suggested absence of genotoxicity and clonal dominance caused by the AAV donor integration.

## Discussion

In this work, we outline essential preclinical studies aimed at progressing an HDR-based genome editing strategy to treat WAS toward an upcoming phase I/II clinical trial. The path from proof-of-concept to clinical application of advanced therapeutic medicinal products (ATMPs) based on the CRISPR/Cas system is complex and currently lacking standardized guidelines from regulatory agencies, resulting in great variability in practices across studies, knowledge fragmentation and increased research costs, thus posing significant barriers in reaching patients. Our study provides a potential framework, demonstrating one approach to conducting preclinical studies for HSPC-based gene editing strategies in accordance with the latest regulatory guidance on this matter.

Our recent proof-of-concept study provided specificity, toxicity and efficacy data supportive of continued development of genome editing to treat WAS (Rai et al. 2020). The platform relies on CRISPR/Cas9 reagents targeting the start codon of the *WAS* gene and an AAV6 with a codon optimized WAS cassette flanked by WAS homology arms which serves as a template for HDR-mediated repair. We demonstrated that the delivery of editing reagents to WAS HSPCs led to the integration of the therapeutic cassette in the *WAS* locus and to full correction of all functional defects in B- and T-cells, macrophages and platelets both *in vitro* and *in vivo* upon HSPC transplantation. This was achieved through the integration of the corrective gene downstream of the endogenous WAS regulatory elements, enabling regulated expression of the WAS protein across all lineages at levels comparable to those in healthy cells. This represents a significant advantage over existing lentiviral-based approaches and underscores the value of site-specific integration achieved through HDR strategies. In this study, we focussed on key aspects essential for successful clinical translation, including: 1) the ability to manufacture this ATMP at clinical scale while preserving HSPC fitness and BM repopulating capacity, and 2) the overall safety of the gene editing procedure. The former represents a major challenge in the HDR field, limiting the broader application of these technologies to the treatment of blood disorders that require high levels of donor chimerism following transplantation. Many preclinical reports have indeed highlighted that animals transplanted with *ex-vivo* manipulated HSPCs undergoing HDR display an overall lower human engraftment rate than those infused with control cells (Lee et al. 2024). Similarly, a clinical trial (NCT04819841) utilizing CRISPR/Cas9 and an AAV6-based HDR template to introduce a functional copy of the β-globin gene for Sickle Cell Disease (SCD) (Lattanzi et al. 2021) showed promising preclinical results but unsuccessful clinical outcome, as the treated patient experienced incomplete hematopoietic recovery, likely due to cell toxicity associated with the editing process. Evidence indicates that while the frequency of corrected cells in vivo remains comparable to input levels soon after transplantation, it markedly diminishes over time – highlighting that primitive long-term HSCs are likely minimally corrected, underrepresented in the transplant product, or impaired in their engraftment and self-renewal ability post-manipulation (Ferrari et al. 2020; Lee et al. 2024; Maganti et al. 2021). Therefore, thorough investigation on the quality of the manufacturing product is mandatory for the clinical success of the gene editing strategy proposed. In this study, we established a clinically-ready protocol for gene editing that ensures preservation of key quality features of HSPCs when manipulated at large scale. Considering that for a successful autologous stem cell transplant it is desirable to infuse 2.5–6 × 10⁶/kg (EBMT guidelines) (Carreras 2019), and that higher doses (5-10 × 10⁶/kg or more) have been associated with better engraftment and reduced complications (Scheid et al. 1999; Tucci et al. 2019), we edited an average of 110 × 10⁶ cells per each donor source used to mimic a possible manufacturing scenario. By using GMP-grade reagents and optimised culture and electroporation procedures, we were able to achieve HDR rates comparable to those observed when editing at medium and small scales, while attaining viability rates comparable to unmanipulated cells both in the fresh and in the cryopreserved product. Most importantly, large-scale edited cells and control cells generated comparable numbers of colonies in solid cultures and engrafted the bone marrow of recipient mice at similar frequencies, highlighting the preservation of HSPC repopulating capacity despite ex vivo genetic manipulation. Overall, all 4 manufacturing runs met or exceeded the Batch Release Criteria set out by our GMP unit at the Great Ormond Street Hospital NHS Foundation Trust (UK) for the release of gene therapy-based ATMPs, including cell purity, viability, editing frequency and colony-forming capacity, confirming the robustness and suitability of the devised protocol. Notably, fine tuning of the electroporation procedure to deliver the CRISPR/Cas RNP complex yielded minimal cell toxicity as highlighted by all QC tests and confirmed by RNA-seq analysis and engraftment rates *in vivo*, representing a significant improvement compared to other reports (Schmiderer et al. 2020; Vavassori et al. 2023)

A key limitation of our platform is the consistent decline in the percentage of gene-corrected cells following engraftment, a challenge commonly observed in approaches that rely on harnessing the HDR pathway for HSPC correction, as previously discussed (Lee, Lozano, and Dunbar 2021). This defect is likely caused by activation of a DNA damage response and senescence-like cell phenotype mediated by p53- and p38 MAPK-ROS signalling pathways (Della Volpe et al. 2024) (Schiroli et al. 2019). RNA-seq analysis performed on CD34+ cells edited at large scale compared to unmanipulated controls confirmed the presence of gene expression dysregulation pointing to disruption of cellular pathways linked to apoptosis, cell cycle arrest and inflammation in our edited products. Because the use of an AAV donor molecule has been indicated as one of the main culprit (Ferrari et al. 2022; Schiroli et al. 2019) and because partial reversion of the phenotype and preservation of HSPC function has been demonstrated when pharmacological intervention targeting these pathways has been employed, we further optimized our ex vivo gene editing protocols by reducing the AAV burden and by transiently inhibiting the p53 pathway. The addition of i53 mRNA alone improved outcomes, enhancing colony-forming capacity in semisolid cultures and significantly increasing the number of engrafted, gene-corrected cells in hematopoietic organs. However, the combination of 53BP1 inhibition with a tenfold reduction in AAV dose yielded even greater benefits, resulting in up to a fivefold increase in the frequency of engrafted CD34⁺ cells carrying the WAS cassette knock-in. While the results may be skewed by the oligoclonal nature of the mouse model used and as such the definitive ability of edited cells to reconstitute a healthy hemopoiesis post transplantation will only be determined in patients, this piece of data emphasises the need of careful fine tuning of manufacturing protocols to achieve therapeutic outcomes, while shedding light on a promising route to preserve HSPC function —paving the way for more effective CRISPR therapies.

Despite the efficiency and versatility of the CRISPR/Cas system, unintended genetic modifications at off-target sites remain a major safety challenge. These off-target edits, which may involve the generation of indels, have the potential to interfere with gene expression and regulation, sometimes leading to harmful consequences such as oncogenic changes. Additionally, genome instability can arise from larger structural alterations at both intended and unintended sites, which can ultimately result in chromosomal loss (Boutin et al. 2022; Lopes and Prasad 2023). Therefore, comprehensive profiling of off targeting events is needed before reaching clinical translation. Despite the development of a large plethora of methodologies to detect such genomic edits, no single method can exhaustively characterize the off-target spectrum in the cell product (Cromer et al. 2023) and as such the use of orthogonal techniques has been recommended by the Food and Drug Administration. In this study, we evaluated genotoxicity in a comprehensive way by combining in silico prediction with a genome-wide, unbiased GUIDE-seq method to detect off-targeting, and complemented this with genome stability assessment through CAST-seq, karyotyping and analysis of AAV donor vector integration. Off-target detection conducted in edited CD34+ cells from 3 WAS donors and 4 healthy donors revealed an overall safe profile for the selected gRNA targeting *WAS,* but highlighted the need of a personalised approach that takes in consideration genomic variance across individuals (Cancellieri et al. 2023; Scott and Zhang 2017). Indeed, 1 WAS and 1 healthy donor out of 7 showed significant cutting at the fifth intron of *COL23A1* in the product examined in vitro, with one of them showing emergence of the same pattern 12 weeks after transplantation in one mouse. The same holds true for the CAST-seq analysis, where a low frequency translocation between the on-target and a natural break site on chromosome17 was detected only in mice transplanted with cells derived from one specific healthy donor, but not in the cryopreserved product before transplantation. This observation also underscores the importance of evaluating the progression and functional implication of a CRISPR/Cas-induced mutation when detected in the manufactured product, which was tackled in this study by an orthogonal and longitudinal analysis at different time points. Indeed, most off-target mutations tend to arise in intronic or other non-coding regions, where they generally pose little risk to cellular function (Smith et al. 2020). When mutations do occur in exonic regions, they are often loss-of-function variants that are naturally eliminated through negative selection, making their persistence in the patient unlikely. In contrast, gain-of-function mutations are more worrisome, as they may confer a growth advantage that drives clonal expansion and potentially contributes to cancer development. In our case, the intronic off target site detected in one scale-up product was lost over the 10-day culture time in vitro, suggesting negative selection. However, the same off target site was found edited in the BM of one out of 12 examined mice 12 weeks post-transplantation, indicating that despite its low frequency, the mutation was harboured by a fraction of BM-repopulating CD34+ cells and was maintained in an *in vivo* setting. This underscores the critical need for patient monitoring following transplantation, as further emphasized by the detection of a de novo chromosomal rearrangement — between the on-target site and a natural DNA break site — observed only after cell engraftment in mice. This rearrangement was likely driven by the proliferative stress placed on a limited pool of edited HSPCs during hematopoietic reconstitution (Flach et al. 2014), a phenomenon that might be prevalent in many gene editing clinical trials where a limited number of corrected HSPCs is able to engraft. Whether these undesired genomic aberrations will lead to clonal dominance and tumorigenesis in the patients, it will only be determined with long-term patient surveillance post transplantation. Nevertheless, in our 12 weeks-long in vivo experiments, we could not detect any sign of cell and tissue toxicity, as evidenced by the wellbeing of transplanted mice and absence of lesions in the hematopoietic organs analysed by immunohistochemistry. Tracking of indels variants at the *WAS* on target locus before and after transplantation confirmed absence of skewed proliferation of cells bearing specific mutations at the genomic site, indicating that infusion of edited CD34+ cells that underwent NHEJ-rather than HDR-mediated correction of generated DSBs is likely not harmful to the patient; this observation was also confirmed by counterselection of indels-bearing CD34+ cells *in vivo*. Finally, our comprehensive preclinical study contributed to the characterization of the impact of AAV integration to HSPC engraftment potential and overall genotoxicity, an aspect that has been largely overlooked by the HDR field. Although detection was limited only to a fraction of cells bearing integration of the donor molecule, tracking of AAV integration before and after transplantation further highlighted absence of clonal selection and abnormal expansion of edited CD34+ cells in vivo, with a polyclonal integration pattern and no integrations detected into or nearby oncogenes.

In summary, we have reported the establishment of a clinically scalable and regulatory-compliant CRISPR/Cas9-AAV6 HDR gene editing platform for diseases that could be treated by ex vivo HSPC gene editing, demonstrating functional correction of HSPCs and emphasizing the importance of optimized manufacturing protocols and thorough genotoxicity assessments to ensure efficacy, safety, and long-term engraftment potential. The study outlines a structured, regulatory-aligned approach for translating CRISPR/Cas9-AAV6 HDR strategies into clinical trials, and sets a precedent for regulatory-grade product characterization, potentially serving as a model for other gene-editing therapies.

## MATERIALS AND METHODS

### Study design

The aim of this preclinical study was to assess efficacy, safety, and clinical-scale manufacturing feasibility of the manufactured product which uses ex vivo Cas9-RNP coupled to rAAV6 mediated coWAS donor cassette delivery to correct mutations in the WAS gene in human HSPCs. Our main goal was to comprehensively determine the safety of our gene editing platform to translate it into the next generation of therapeutic tools for WAS.

### Ethics and animal approval

For usage of human CD34+ HSPC from healthy and WAS donors, informed written consent was obtained in accordance with the Declaration of Helsinki and ethical approval from the Great Ormond Street Hospital for Children NHS Foundation Trust and the Institute of Child Health Research Ethics (08/H0713/87).

For experiments involving animals, NOD-SCID-IL2Rg^-/-^ (NSG) mice were purchased from Charles River and maintained in specific-pathogen-free (SPF) conditions. Mice were housed in a 12-h day-12h-night cycle with controlled temperature and humidity. The ventilated cages had sterile bedding and everyday supply of sterile food and water in the animal barrier facility at University College London. The procedures involving mice were designed and performed in accordance with UK Home Office regulations, and experiments were conducted after approval by the University College London Animal Welfare and Ethical Review Body (project license 70/8241).

### AAV6 vector design, production, and purification

GMP compliant therapeutic AAV6 vector plasmid was generated by cloning codon-optimised 1.5-kb-long WAS cDNA (coWAS) flanked by 405bp long left and right homology arms described previously (Rai et al) into the pBB4-MCS plasmid (Dr. Leszek Lisowski, Univ of Sydney), with inverted terminal repeats derived from AAV2.

AAV6 vectors were either purchased from Vigene Biosciences or produced in house as described previously (Rai et al). Briefly, producer AAVPro293T cells (Takara Bio; Cat# 632273) were thawed at 37°C and expanded in Dulbecco’s modified Eagle’s medium (DMEM) supplemented with 10% FBS and 1% penicillin/streptomycin. The cells were expanded and plated in 10cm plates a day before transfection. 24 hours later, producer cells were cotransfected with the donor template plasmid (described above) and pDGM6 Rep/Cap helper plasmid, using a polyethylenimine-based transfection method with a change of medium performed the next day. 72 hours after transfection, the resulting cell lysates and purified media were purified using iodixanol gradients-based ultracentrifugation. Viral genomic titer was determinated by ddPCR using ITR specific primers and probes as listed in Supp Table 2. The final coWAS_AAV6 preps had an average titer of 1 × 10^13^ vg/ml which were subsequently aliquoted and stored at −80°C until use.

### Isolation and in vitro culture of CD34+ HSPCs

Human HSPCs were isolated by the immunomagnetic selection of CD34+ cells from G-CSF mobilized apheresis products from healthy donors (Caltag Medsystems and All cells). Before CD34+ selection, the apheresis product was washed with CliniMACS PBS/EDTA buffer containing 0.5% (v/v) human serum albumin to remove platelets. CD34+ HSPCs were then either selected manually (SU1&2) or isolated by positive selection on the CliniMACS Plus instrument (Miltenyi Biotec, Bergisch Gladbach, Germany) for SU3&4. Isolated HSPCs were either freshly plated in culture (SU2-4) or cryopreserved in Cryostor freezing medium (CS5; Stemcell) (SU-1) and thawed at the time of experiment. HSPCs were cultured at a cell density of 0.5 × 10^6^/ml in StemSpan Serum-Free Expansion Medium II (STEMCELL Technologies, Vancouver, Canada) supplemented with human cytokines: stem cell factor (SCF, 100 ng/ml), thrombopoietin (TPO, 100 ng/ml), Fms-like tyrosine kinase 3 ligand (Flt3-L, 100 ng/ml), Interleukin-6 (IL-6, 100 ng/ml), IL-3 (30 ng/ml) and penicillin-streptomycin (20 U/ml), stemregenin1 (0.75M; STEMCell Technologies, Vancouver, Canada) and UM171 or UM729 (35 nM; STEMCell Technologies, Vancouver, Canada).

### Gene correction procedure and associated analyses

GMP complaint synthetic sgRNA containing 2-O-methyl-3′-phosphorothioate at the three terminal positions at both 5′ and 3′ ends targeting *WAS* locus was provided by synthego. The *WAS* sgRNA was >99% pure and the target sequence for is 5′-GCAGAAAGCACCATGAGTGGGGG -3′. GMP SpyFi Cas9 nuclease was purchased from Aldevron (Fargo, ND, USA). CD34+ HSPCs were either thawed or cultured from fresh apheresis in the Stemspan SFEMII medium supplemented with early active cytokine cocktail (SCF, TPO, Flt3-L, IL6 and IL3) alongwith UM171 and Stemregenin-1 for a period of 48 hours. RNP electroporation was performed at day2 using the Maxcyte electroporator. Briefly, the cells were collected, counted, and spun at 300g for 5min, following which the supernatant was discarded carefully and the cell pellet was washed with pre-warmed 1ml of Maxcyte electroporation buffer (HyClone). Large scale RNP electroporations were performed using a CL1.1 cartridge (suitable for 100-700M cells) in a total volume of 1.8ml and with an OC-100 cartridge in a total volume of 100ul for medium scale (program HSC-3). Electroporated cells were then plated at a cell density of 1.25 × 10^6^ cells/ml in the cytokine-supplemented medium as above. After 15 minutes, rAAV6 carrying the coWAS cassette was dispensed onto cells at multiplicities of infection of 2.5 × 10^4^ vector genomes per cell based on titers determined by ddPCR and incubated for 24 hours. After 24 hrs, a medium exchange was performed to dilute or remove any residual coWAS-AAV6, and the CD34+ HSPCs were cultured for an additional 12 to 24 hours before cryopreservation. Additionally, a subset of cells from the three groups were cultured for an additional 2-3 weeks for in vitro genotoxicity analysis.

### Methylcellulose CFU assessment

At day 1 post editing, the HSPCs from the transduced and the control groups were plated into 12 well plates containing MethoCult (STEMCELL Technologies, Vancouver, Canada). Two weeks later, colonies were appropriately scored based on morphological appearance in a blinded manner. Subsequently, colonies were harvested and lysed for genomic DNA extraction followed by ddPCR analysis for targeted integration of WAS cassette.

### Targeted integration analysis by digital droplet PCR

For the measurement of targeted integration of codon-optimised WAS cDNA in the edited samples, ddPCR was performed as previously described (Rai et al., 2020). Briefly, genomic DNA was extracted using DNeasy blood and tissue kit (Qiagen; Cat# 69504) and a total of 60ng was mixed with 10 μM each of target primer and FAM probe mix, 10 μM each of reference primer and HEX probe mix, 1× ddPCR Supermix probe without dUTP (Bio-Rad, UK) and nuclease-free water. The primers and probes sequences are detailed in Supplementary Table 2. The individual droplets were generated using QX100 Droplet Generator (Bio-Rad) and subsequently amplified in a Bio-Rad PCR thermocycler. The optimised amplification steps were: step 1 – 95 °C for 10 min; step 2 (49 cycles) – 94 °C for 1 min, 60 °C for 30 s, 72 °C for 2 min; and step 3 – 98 °C for 10 min. The Droplet Reader and QuantaSoft Software (both from Bio-Rad) were used to record and analyse the positive and negative fluorescence droplets according to the manufacturer’s guidelines. The percentage of integration was calculated as the ratio of FAM to HEX signal after normalisation against the reference signal.

### Karyotype Analysis

CD34+ HSPCs from healthy mobilised peripheral blood were electroporated as described above. After 2-4 days in culture, 3-5× 10^6^ cells from edited and transduced group as well as WT control cells were incubated overnight with 0.6ug/ml of mitotic inhibitor Colcemid (KaryoMAX Colcemid Solution in PBS Life Technologies, Carlsbad, CA), thereafter incubated in hypotonic solution (0.075M Potassium Chloride (KCl) added dropwise) at 37C for 25 minutes, followed by the dropwise addition of the fixative (3:1 methanol/acetic acid). Karyotype analysis was then outsourced to Cambridge Biosciences. The metaphase spreads were aged at room temperature for 5 days and G-banding were performed following standard methods with a few modifications: slides were incubated in Sorensen buffer at room temperature for 2 minutes, followed by 2 minutes in Wright stain solution (0.5ml Wright stain and 1.5 mL of Sorensen buffer). A total of 270 metaphase spreads from three independent HSPC donors and editing experiments were analyzed with a Leica DM-2500 Bright Field Microscope and the software used to analyse is Captured images were analysed using the software Cytovision version 7.7.

### Genome edited CD34+ HSPC xenotransplantation studies in NSG mice

Edited cells as well as WT and RNP controls frozen 36-48 hours post editing as detailed above were thawed and 5 × 10^6^ viable cells were injected intravenously into each NSG mouse after sublethal irradiation (2Gray) using a 27 gauge × 0.5-inch needle. 8 weeks post transplantation, mice peripheral blood from tail vein was lysed with 1× RBC lysis buffer (ThermoFisher Scientific) and stained with anti-human CD45 APC antibody (clone HI30, BioLegend) to evaluate human CD45+ engraftment by FACS. At week 14, post transplantation, level of human engraftment and lineage composition were determined in the bone marrow, peripheral blood, spleen and thymus. Briefly, bone marrow cells were harvested by flushing tibiae and femurs with 1× PBS and passing through 40 μm strainer. Mononuclear cells were blocked with Fc blocking solution (Bio-Legend) and stained with the following anti-human antibodies: CD45 PerCP Cy5.5 (clone HI30, Biolegend), CD19 APC Cy7 (clone HIB19, BioLegend), CD33 PECy7 (clone 67.6, Biolegend) and CD3 APC (clone OKT3, BioLegend) for FACS analysis. To determine human stem cell composition within bone marrow, cells were stained with the following antibody panel: CD34 BV421 (clone 561, Biologend), CD45 PerCP/Cy5.5 (clone HI30, Biolegend), CD38 APC-Cy7 (clone HIT2, Biolegend), CD90 PE-Cy7 (clone 5E10, Biolegend). Spleens and thymi were grinded against 40 μm strainer and stained for the lineage antibody panel and CD19 APC Cy7 and CD3 APC Abs respectively.

Error! Reference source not found.

Histopathological analysis on bone marrow smears and spleen sections obtained from transplanted NSG mice was carried out by Histologix in GLP conditions. Spleen samples were fixed in 10% NBF for 24-48 before being transferred to 70% ethanol. For sectioning, 5µm sections were taken for each spleen sample, picked up on Snowcoat® slides and dried at 37°C overnight. The spleen tissues were trimmed to standardised trimming planes, placed in tissue cassettes and processed to paraffin wax and sectioned. The slides were stained with Haematoxylin and Eosin (H&E) and digitally scanned at x20 magnification using a Hamamatsu NanoZoomer v2. The bone marrow smear preparations were sprayed fixed on to the slide using CytoFix pump spray (CellPath Ltd, UK) and were stained with Haematoxylin and Eosin (H&E) for pathology review and scanning as mentioned above.

### Off-target analyses

Frozen male peripheral blood derived CD34+ HSPCs were cultured for 2 days prior to electroporation with RNP complex as described above. Day 4&14 post RNP electroporation, genomic DNA was extracted and PCR was performed using off target primers for the combined sites retrieved from COSMID and Guide-Seq (Rai et al 2020). PCR amplicons of WT cells were taken as negative untreated control. Forward and reverse off target primers were designed around the predicted off-target sites in order to amplify fragments of ∼200-300 bp (Supp table 2). The PCR-purified amplicons were subjected to end repair, adaptor ligation and an indexing PCR using the NEBNext® Ultra™ II DNA library prep kit for Illumina (New England BioLabs) according to the manufacturer’s instructions. The denatured amplicons were loaded at 12 pM into the Illumina MiSeq Reagent Kit V2 - 500 cycle according to the manufacturer’s instruction. The FASTQ files were analysed for indels using the command line version of CRISPResso49 considering 40 bp around the supposed cleavage site and not considering substitutions to reduce background signals. The derived INDEL proportions of the treated samples were statistically compared to the corresponding untreated values in a one-tailed Z-test. For this purpose, the RNP treated values were corrected by the standard deviation of the untreated samples to account for the variability of the measurements.

### Detection of translocations: chromosomal aberrations analysis by single targeted linker-mediated PCR sequencing (CAST-Seq)

CAST-seq was performed as described in “Turchiano et al. 2021”. Genomic DNA was harvested from the sample before and after xenotransplant, fragmented and linker ligated using the NEB Next Ultra II FS DNA Library Prep Kit and NEB Next Ultra II Ligation Master Mix and Ligation Enhancer (New England Biolabs). Specific primers were designed for the WAS locus: WAS_Bait: TCCAAGCATCTCAAAGAGTC and WAS_Decoy: GCAGGGTAAGAAGAGGAAC for the first PCR; and WAS_Nested: GACTGGAGTTCAGACGTGTGCTCTTCCGATCTGGAGGGTATGTTCTGCTGAACC for the second PCR. For CAST-seq of the ex vivo samples, 500ng of gDNA was used as starting material, while for the samples post-transplantation 20-100ng were used. The PCR reactions were purified and barcoded with NEB Next Multiplex Oligos from Illumina (New England Biolabs). AMPure XP beads purification (Beckman Coulter) was set at 0.8X volume to select fragments >180bp. The libraries were sequenced through the Illumina MiSeq platform and analysed with the CAST-seq pipeline (https://github.com/AG-Boerries/CAST-Seq) calling as true mutation events sites that were significantly enriched with respect to the UT control and present in at least two replicates.

### Indel deep sequencing analysis for mutational tracking

Off-target sites identified with the CAST-seq, GUIDE-seq and COSMID tool were subjected to direct PCR amplification and deep sequencing. Amplicons of ca. 300bp were barcoded using NEBNext_Ultra II DNA library prep kit for Illumina and sequenced through the Illumina MiSeq platform. The sequencing adapters were trimmed using cutadapt (Martin 2011) using the parameters -j 5 -m 20 -u −1 -U −1. The reads were subsequently analysed using the CRISPResso2 pipeline (Clement et al. 2019), considering an indel window of 10 bp around the putative cleavage site and disregarding substitutions (parameters: --quantification_window_size 10 --ignore_substitutions). The indel proportions obtained from the edited samples were compared with their relative untreated control by means of the one-tailed Z-test corrected by the standard deviation from the untreated control proportions to account for measurement variability. The mutation tracking analysis was derived by parsing the variant frequency data generated by CRISPResso2 and intersecting the mutation across samples with R package. Any variant which was marked as “unedited” by CRISPResso2 was considered wild-type for the target site and any variant not detected in a particular sample was given a value of zero. For visualisation, the frequency of each variant from sample WT was subtracted from the frequency of that variant in every other sample. The selection of the top 10 up- and down-represented variants was based on the difference between sample WT and the mouse derived samples. Heatmaps were generated with the pheatmap package (Kolde 2019).

### Integration site (IS) distribution analysis

To assess AAV integration, we adopted a sonication-based linker-mediated PCR method (SLiM), as previously described (Calabria et al. 2023). Briefly, genomic DNA was sheared using a Covaris E220 Ultrasonicator (Covaris Inc) (Cesana et al. 2021; Gentner et al. 2021), generating fragments with a target size of 1000 bp. The fragmented DNA was subjected to end repair, 3′ adenylation, and ligation (NEBNext Ultra DNA Library Prep Kit for Illumina, New England Biolabs) to custom linker cassettes (LCs; Integrated DNA Technologies). LC sequences contained an 8-nucleotide barcode for sample identification. Ligation products were subjected to 35 cycles of exponential PCR with primers (available upon request) complementary to different regions of the AAV genomes (supplemental Figure 2A; available on the Blood website) and to the LC. For each set of AAV-specific primers, the procedure was performed using ∼50 to 100 ng of sheared DNA. Then, 10 additional PCR cycles were run to include the sequences required for sequencing and a second 8-nucleotide DNA barcode. PCR products were quantified via qPCR using the Kapa Biosystems Library Quantification Kit for Illumina, following the manufacturer’s instructions. qPCR was performed in triplicates on each PCR product diluted 10:3, and the concentrations were calculated by plotting the average cycle threshold values against the provided standard curve. Finally, the amplification products were sequenced using Illumina Next/Novaseq platforms (Illumina). After sequencing, a dedicated bioinformatics pipeline, recombinant adeno-associated vector integration analysis (RAAVioli), was used to analyze the amplified sequences for integration site identification.

### Statistical analysis

All statistical tests on experimental groups were done using Prism7 GraphPad Software. The exact statistical tests used for each comparison are noted in the figure legends. For making multiple comparisons, one-way ANOVA Kruskal-Wallis multiple comparisons test with one variable and two-way ANOVA with Bonferroni post-test in case of two variables. For comparing the average mean of two sample groups, we used the unpaired Student’s *t* test to reject the null hypothesis (*P* < 0.05).

## Data availability statement

The data supporting the findings of this study are available within the article and its supplementary materials. Additional data or materials, including protocols and reagents, are available from the corresponding author upon reasonable request.

## Acknowledgements

We would like to thank all past and current members of the Cavazza and Santilli groups for their contribution in conceptualizing this study and experimental support. We thank Dr. Ayad Eddaoudi and his team (Flow Cytometry Core Facility, University College London) for assistance with flow cytometry and the team at UCL Genomics Core Facility, University College London for assistance with NGS sequencing. This research, A.C. and A.N. were supported by the GOSH Children’s Charity-LifeArc Therapeutic Accelerator Grant (VS0420) and the Sparks-GOSH Children’s Charity Grant (V4522). This research was funded, in part, also by the Wellcome Trust [217112/Z/19/Z]. T.W. was supported by the Jeffrey Modell Foundation. G.S., A.J.T., and A.C. were also supported by the NIHR Biomedical Research Center at the Great Ormond Street Hospital for Children NHS Foundation Trust and University College London.

## Author contributions

A.N. designed and performed experiments, analysed and illustrated data. W.V., T.W, F.Z., R.G., E.G.C., C.C., G.T., D.C. performed experiments and analysed data. H.A. contributed to the design of experiments and illustrated data. E.M., A.J.T, G.S., D.C., G.T. and A.C. contributed to the design and conceptualization of the study or parts of it. A.C. designed and supervised the study and the experiments, analysed data, provided funding, and wrote the manuscript. All authors reviewed the manuscript.

