## Supplemental data for "In-Depth Characterization of Stem Cell Potency and Genotoxicity for Clinical-Scale Ex Vivo CRISPR/Cas9 Gene Editing"

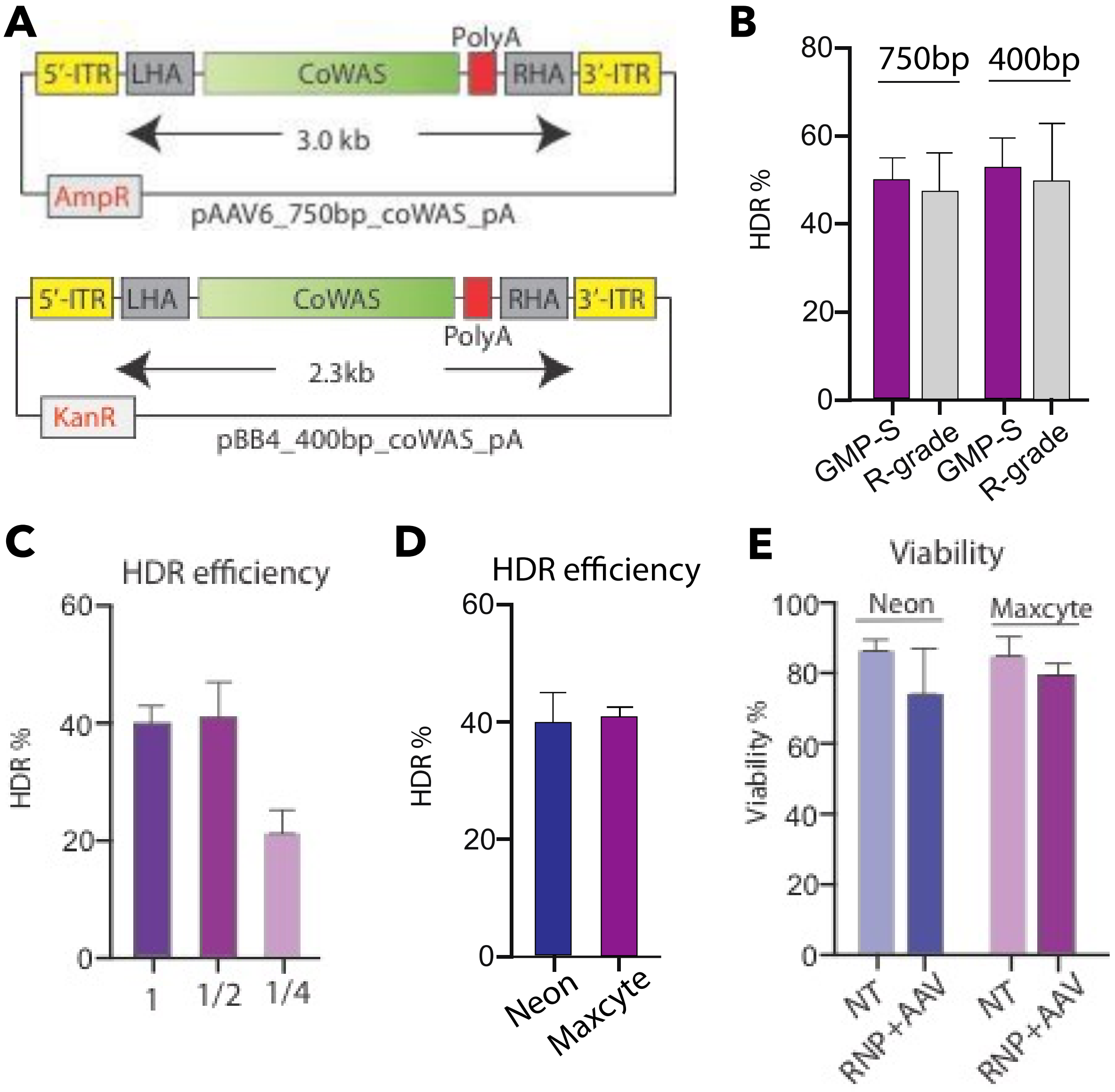

**Figure S1. HSPC gene editing scale-up protocol and reagents optimization.**
**(A)** maps of the AAV donor molecules used in the proof-of concept study by Rai et al[1] (pAAV6_750bp_coWAS_pA) and in this study (pBB4_400pb_coWAS_pA). (B) Frequency of coWAS cassette knock-in via HDR in HSPCs (n=3 biological replicates) when using the two different AAV donor vectors (as in (A)) and cGMP- or research-grade reagents. (C) Frequency of coWAS cassette knock-in via HDR in HSPCs (n=3 biological replicates) when using different RNP doses. (D-E) Frequency of coWAS cassette knock-in via HDR in HSPCs (D) and cell viability (E) when electroporating cells with different electroporation machines (n=3 biological replicates). Unmanipulated HSPCs were used as control.

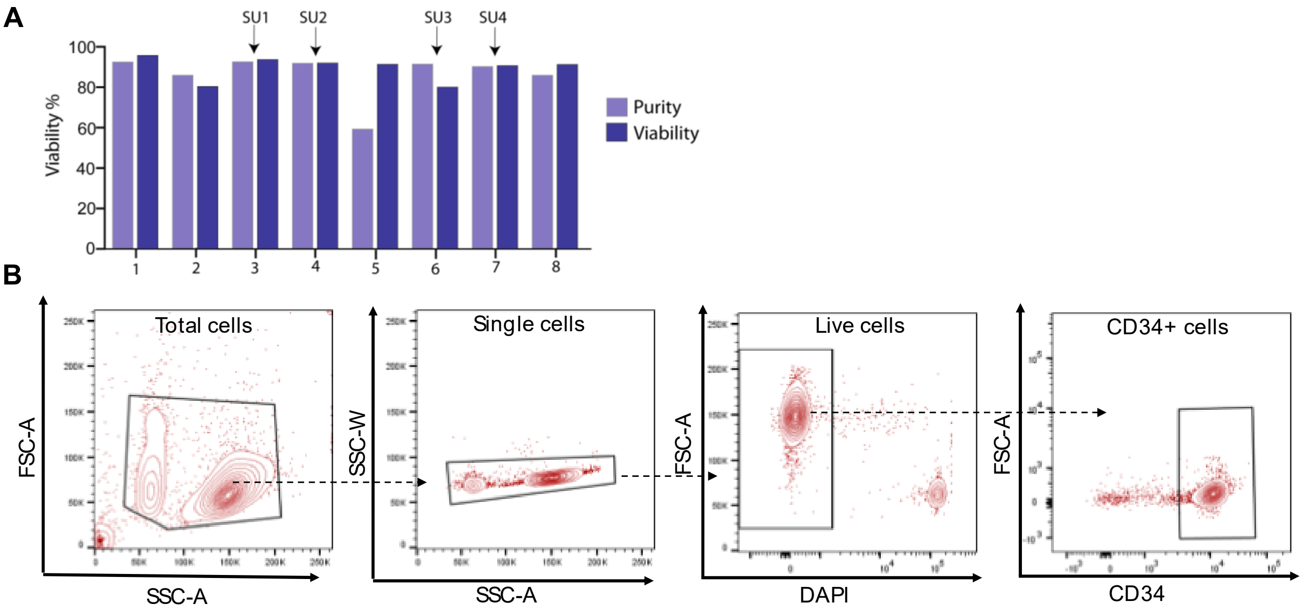

**Fig S2: Purity and viability of HSPCs isolated from leukapheresis products** (A) CD34+ isolation was carried out from G-CSF mobilised leukapheresis from eight healthy donors. Purity (fraction of CD34+ cells) and viability post selection is shown. Arrows indicate the donors used for the 4 scale ups in this study. (B) Gating strategy used to assess CD34+ purity and viability.

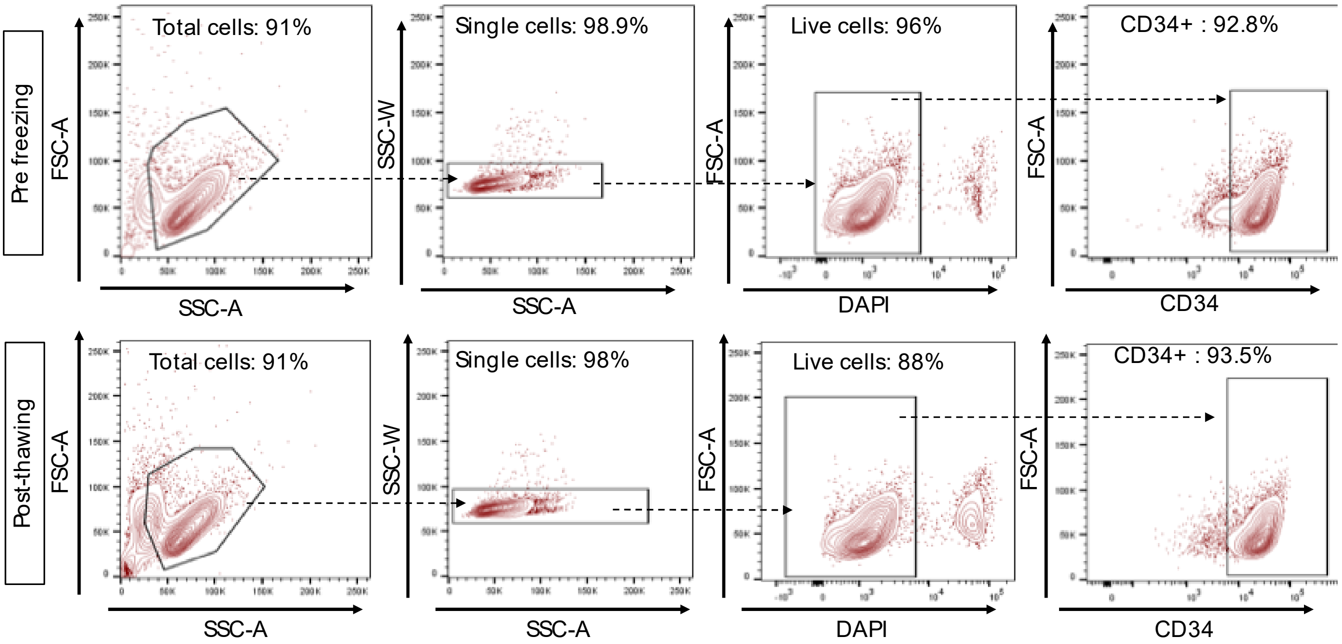

**Fig S3: Gating strategy to assess HSPC purity and viability pre- and post-cryopreservation** (A) CD34+ isolation was carried out from G-CSF mobilised leukapheresis from eight healthy donors. Purity (fraction of CD34+ cells) and viability post selection is shown. Arrows indicate the donors used for the 4 scale ups in this study. (B) Gating strategy used to assess CD34+ purity and viability.

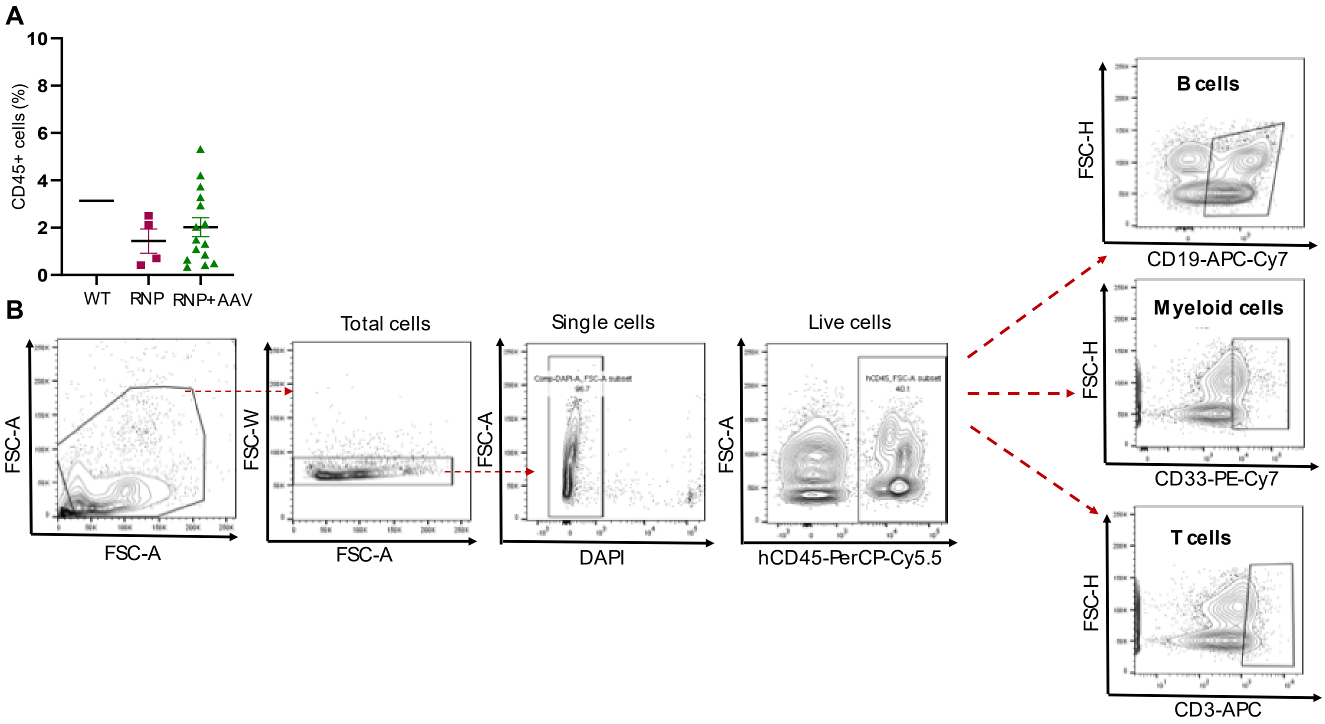

**Fig S4: Assessment of human chimerism in xenotransplanted NSG mice.** (A) Engraftment frequencies in the peripheral blood of tail vein-bled transplanted NSG mice at 8 weeks post injection, expressed as the frequency of CD45+ cells detected by flow cytometry. (B) Representative gating strategy for the analysis of human cell lineages in hematopoietic organs of transplanted mice.

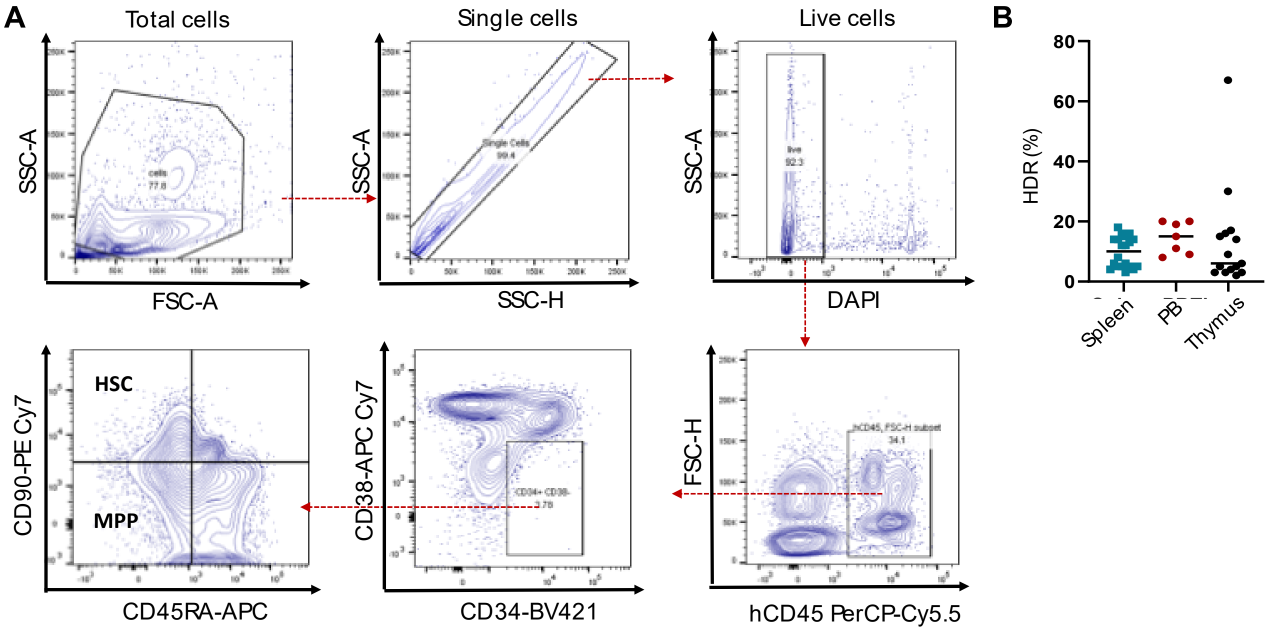

**Figure S5: Assessment of human chimerism and gene editing frequency in xenotransplanted NSG mice**. **(A)** Representative gating strategy for the for the analysis of HSPC subpopulations in the bone marrow of transplanted mice. Human cells (hCD45+) were isolated from the BM of mice 12 weeks post injections. hCD45+ population was phenotyped into primitive HSCs (CD34+ CD38- CD90+ CD45RA-), MPPs (CD34+ CD38- CD90- CD45RA-) and CD38+ committed progenitors (CD34+ CD38+). (B) Frequency of coWAS cassette integration via HDR in the PB, spleen and thymus of mice transplanted with RNP+AAV HSPCs, as detected by ddPCR.

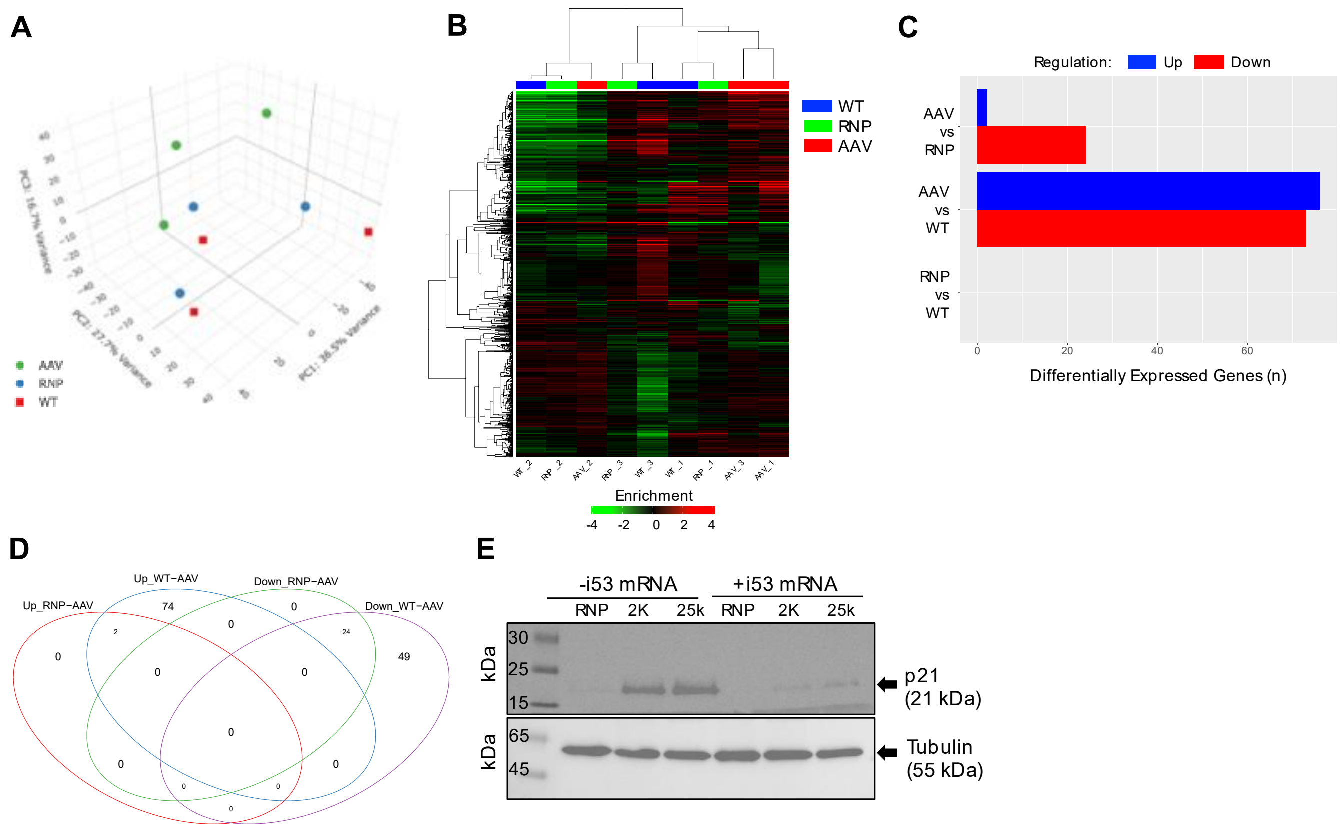

**Figure S6: Dampening of p53 signalling improves engraftment of HSPCs that underwent HDR.** **(A)** Principal Component Analysis conducted on the whole RNA-seq data retrieved from the 3 experimental groups (n=3 biological replicates each). (B) Unsupervised hierarchical clustering of RNA-seq data. (C) Number of genes down- or up-regulated for each pairwise comparison. (D) Venn diagrams displaying the number of differentially expressed genes for each pairwise comparison (RNP vs RNP+AAV or WT vs RNP+AAV) and overlaps among groups. (E) Representative Western blot of p21 expression in HSPCs gene edited using different AAV MOIs and with or without the addition of i53 mRNA. Tubulin was used as a reference for protein quantification.

**A**

| Mouse# | SU1 | | SU2 | | SU3 | | SU4 | |
|---|---|---|---|---|---|---|---|---|
| | BM smear | Spleen | BM smear | Spleen | BM smear | Spleen | BM smear | Spleen |
| WT | WNL | EMH | WNL | EMH | WNL | EMH | WNL | EMH |
| WT | WNL | EMH | WNL | EMH | WNL | EMH | WNL | EMH |
| WT | WNL | EMH | WNL | EMH | WNL | EMH | WNL | EMH |
| WT | WNL | EMH | WNL | EMH | | | | |
| WT | WNL | EMH | | | | | | |
| RNP+AAV | WNL | EMH | WNL | EMH | WNL | EMH | WNL | EMH |
| RNP+AAV | WNL | EMH | WNL | EMH | WNL | EMH | WNL | EMH |
| RNP+AAV | WNL | EMH | WNL | EMH | WNL | EMH | WNL | EMH |
| RNP+AAV | WNL | EMH | WNL | EMH | WNL | EMH | WNL | EMH |
| RNP+AAV | WNL | EMH | WNL | EMH | | | | |
| RNP+AAV | WNL | EMH | WNL | EMH | | | | |
| RNP+AAV | WNL | EMH | WNL | EMH | | | | |
| RNP+AAV | WNL | EMH | WNL | EMH | | | | |
| RNP+AAV | WNL | EMH | WNL | EMH | | | | |
| RNP+AAV | WNL | EMH | WNL | EMH | | | | |
| RNP+AAV | WNL | EMH | WNL | EMH | | | | |
| RNP+AAV | WNL | EMH | WNL | EMH | | | | |
| RNP+AAV | WNL | EMH | WNL | EMH | | | | |
| RNP+AAV | WNL | EMH | WNL | EMH | | | | |
| RNP+AAV | WNL | EMH | WNL | EMH | | | | |
| | normal | minimal | mild | moderate | severe | | | |

**B**

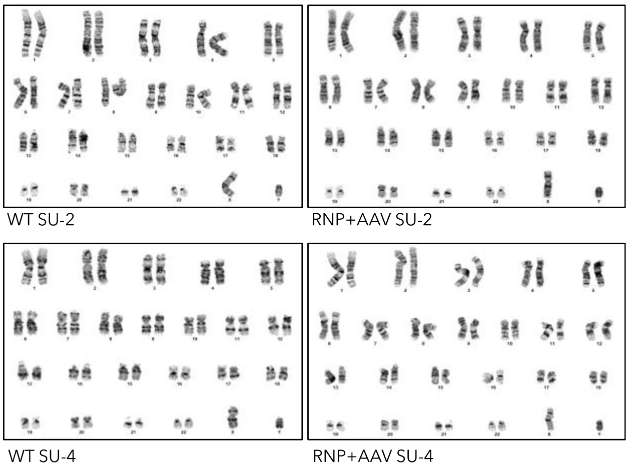

**Figure S7. Evaluating the genotoxicity of the gene editing platform in vitro and in vivo. (A)** Table summarising histopathological results obtained by analysing BM smears and spleen sections upon H&E staining. BM smears revealed expected morphology and normal levels of precursor cells among the groups. Minimal to mild extra medullary haematopoiesis (EMH) was observed in spleen sections, comparable among the groups. (B) Karyotypes of edited (RNP+AAV) and unedited (WT) HSPCs from individual scale up experiments.

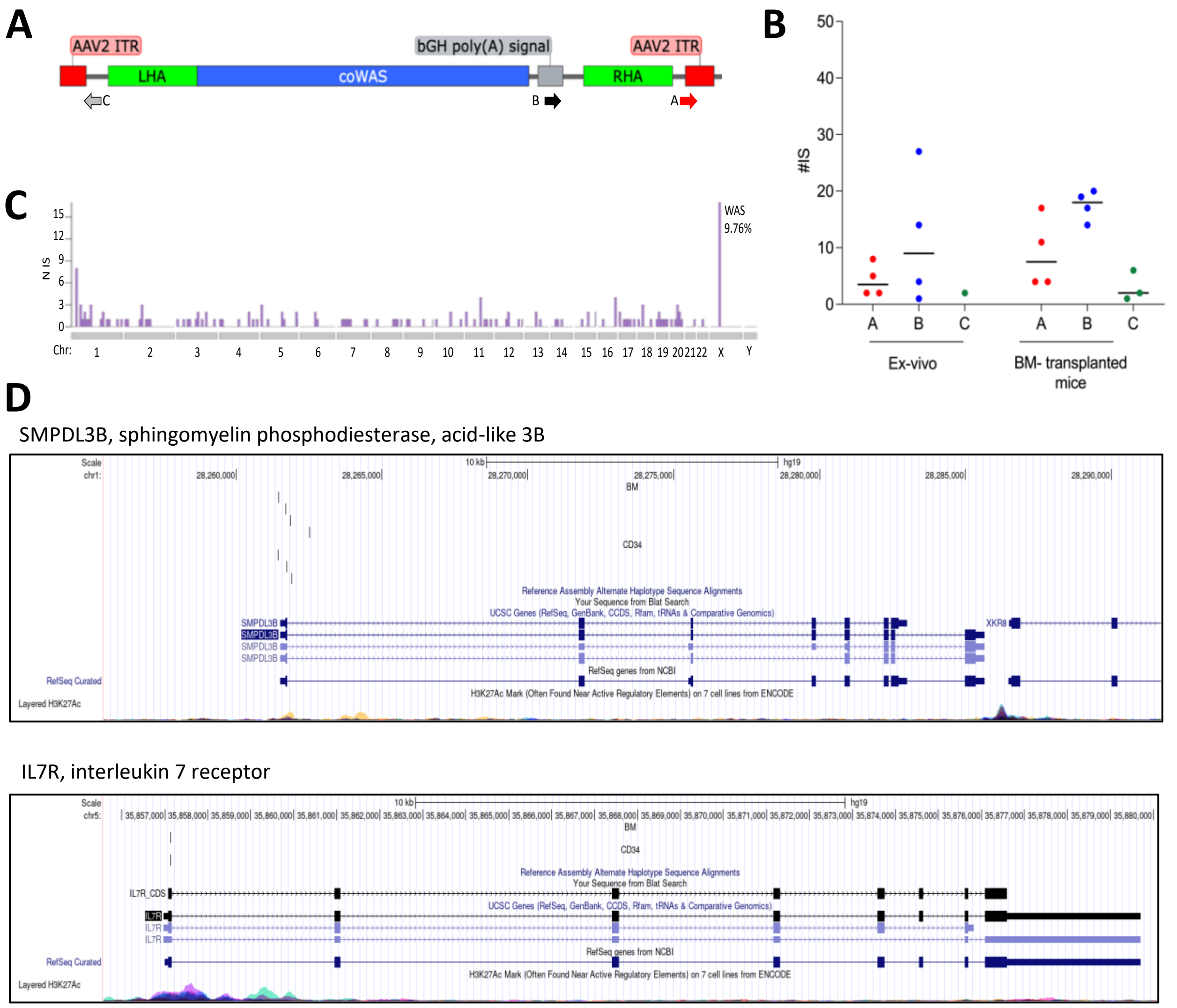

**Figure S8. Analysis of AAV donor molecule integration in gene edited HSPCs.**

**(A)** Schematic representation of the strategy used to map AAV integrations involving different features of the AAV donor vector. (B) Number of AAV integration sites (IS) per each sample analysed, ex vivo and in vivo. (C) Distribution of AAV integrations across chromosomes. High IS integration frequency was observed in chromosome X where the *WAS* locus is located. (D) Representative UCSC Genome Browser snapshot of two genes hit by AAV integration retrieved from edited HSPCs (CD34 track) or from CD45+ cells isolated from the BM of transplanted mice (BM track).

**Table S1. Genomic sites analysed by amplicon deep-sequencing to assess off-target editing.**

| Deetction method | Site name | BED Chromosome | Off-Target start | Off-Target end | Strand | Off-Target Sequence | Mismatches |
|---|---|---|---|---|---|---|---|
| Guide-seq | OT-2 | chr1 | 50322318 | 50322341 | + | GGAAACAG CACCCTGA GAGGAGG | 5 |
| | OT-3 | chr2 | 112824051 | 112824074 | - | GCAGCATG AACAATGG GTGGAGG | 5 |
| | OT-4 | chr2 | 197121373 | 197121396 | + | ACACAAAG CACCATGA GTGGAGG | 2 |
| | OT-5 | chr3 | 2220065 | 2220088 | - | GGAGAAAA CACTCTGA GTGATGG | 5 |
| | OT-6 | chr6 | 159803440 | 159803463 | - | GCAGAGGG CAGAGTGA GTGGAGG | 5 |
| | OT-7 | chrX | 124413658 | 124413681 | + | GCATAAAG CACAAGGA GAGGTGA | 5 |
| Cosmid+NGS | OT-1 | Chr 5 | 178282798 | 178282820 | + | AAAGGAAG CACCATGA GTGGGGG | 3 |

### Table S2. Sequencing results of the translocation detected by CAST-seq in the BM of SU-3 HSPC transplanted mice

| chr | start | end | read (n) | hits (n) | read (n) ctrl | hits (n) ctrl | group | annotation | gene Chr | Distance To TSS | SYMBOL | Genes within 100kb | Oncogenes within 100kb |
|---|---|---|---|---|---|---|---|---|---|---|---|---|---|
| **chrX** | 48674308 | 48687957 | 16465 | 1974 | 2917 | 488 | OMT | Promoter (<=1kb) | 23 | 0 | WAS | GLOD5 ; RBM3; WDR13 ; SUV39H1; WAS | WAS |
| **chr17** | 19697297 | 19697797 | 257 | 34 | 3 | 3 | NBS | Intron (ENST00000325411.9/146802, intron 13 of 16) | 17 | 21182 | SLC47A2 | SLC47A2; ALDH3A1; ALDH3A2 ; ULK2 | |

**Table S3. List and description of AAV integration sites retrieved in this study.**

For each IS matrix there are the following columns:
chr: chromosome number
integration_locus: genomic position of the integration sites
GeneID by RefSeq RefSeq Gene Symbol of the closest gene targeted by vector insertion
Gene strand: genomic orientation of the targeted gene
MhOR/ins: Micro-Homology Region (MhOR): Nucleotides overlap between alignments of AAV and target genome (indicated as negative number) / insertion (ins): De novo insertion of nucleotides between AAV and host genome breakpoint (indicated as positive number)
Junction: nucleotide position on the AAV genome at the vector/cellular genome breakpoint
vector_fragments: number of AAV fragments identified within the vector portion of the reads
sample_columns: column name contained the sample ID, the analyzed tissue, the PCR-set from which the IS was identified and eventually the vector dose. Within the column is indicated the number of different genomic fragments identified for each IS

| Integration sites retrieved in Ex-vivo CD34 and in BM derived cells from transplanted mice | | | | | | | | | | | | | | | |
|---|---|---|---|---|---|---|---|---|---|---|---|---|---|---|---|
| | | | | | | | | Ex-vivo | | | | BM-mice | | | |
| Chr | Integration_locus | strand | Gene_ID | Gene Strand | Mhor/Ins | junction | Vector_frag | SU1_CD34 | SU2_CD34 | SU3_CD34 | SU4_CD34 | SU1_17BM | SU1_21BM | SU3_27BM | SU3_28BM |
| 1 | 21028218 | - | KIF17 | - | -8 | 2419 | 1 | 3 | | | | | | | |
| 1 | 28261437 | + | SMPDL3B | + | -8 | 2337 | 1 | 5 | | | | | | | |
| 1 | 28261444 | + | SMPDL3B | + | -3 | 2290 | 1 | | | | | | 2 | | |
| 1 | 28261703 | + | SMPDL3B | + | -3 | 2378 | 1 | | | | | 2 | | | |
| 1 | 28261741 | + | SMPDL3B | + | -5 | 2305 | 1 | | | 3 | | | | | |
| 1 | 28261862 | + | SMPDL3B | + | -7 | 2346 | 1 | | | | | 2 | | | |
| 1 | 28261890 | + | SMPDL3B | + | -5 | 2321 | 1 | 2 | | | | | | | |
| 1 | 28262505 | + | SMPDL3B | + | -16 | 2337 | 1 | | | | | | 2 | | |
| 1 | 44152858 | + | KDM4A | + | -8 | 2297 | 1 | 3 | | | | | | | |
| 1 | 44152868 | + | KDM4A | + | -18 | 2307 | 1 | | | | 2 | | | | |
| 1 | 44152983 | + | KDM4A | + | -6 | 2275 | 1 | 2 | | | | | | | |
| 1 | 53372169 | - | ECHDC2 | - | -7 | 2322 | 1 | | | | | | | | 3 |
| 1 | 63564339 | + | LINC00466 | - | -1 | 2925 | 1 | | | | 2 | | | | |
| 1 | 66221427 | + | PDE4B | + | 20 | 2909 | 1 | | | | | 2 | | | |
| 1 | 76252309 | - | RABGGTB | + | -4 | 2920 | 1 | | 2 | | | | | | |
| 1 | 89007287 | - | PKN2-AS1 | - | -7 | 2331 | 1 | | | | 2 | | | | |
| 1 | 91852922 | + | HFM1 | - | 4 | 2909 | 1 | | | | | 13 | | | |
| 1 | 93135308 | - | EVI5 | - | -3 | 2919 | 1 | | | | 2 | | | | |
| 1 | 98269320 | + | DPYD | - | 4 | 2909 | 1 | | | | 2 | | | | |

| | | | | | | | | | | | | | | | |
|---|---|---|---|---|---|---|---|---|---|---|---|---|---|---|---|
| 1 | 145103975 | + | SEC22B | + | 0 | 2909 | 1 | | 2 | | | | | | |
| 1 | 153402949 | - | S100A7L2 | - | -4 | 2296 | 1 | | | | 2 | | | | |
| 1 | 153910058 | - | DENND4B | - | -6 | 2300 | 1 | | | | | 2 | | | |
| 1 | 170562747 | + | GORAB | + | 2 | 2910 | 1 | | | | | | 2 | | |
| 1 | 183559474 | - | NCF2 | - | 12 | 2378 | 1 | | | | | 44 | 2 | | |
| 1 | 228463821 | - | OBSCN | + | -16 | 2452 | 1 | | | 2 | | | | | |
| 1 | 240943039 | + | RGS7 | - | -11 | 2278 | 1 | 2 | | | | | | | |
| 2 | 8771152 | + | ID2-AS1 | - | -11 | 2342 | 1 | | | | | | 2 | | |
| 2 | 10895786 | + | ATP6V1C2 | + | -12 | 2448 | 1 | | | 17 | | | | 4 | 2 |
| 2 | 74783270 | + | DOK1 | + | -21 | 2341 | 1 | | | 2 | | | | | |
| 2 | 82534611 | + | DHFRP3 | + | -7 | 2280 | 1 | | | | 2 | | | | |
| 2 | 82534776 | + | DHFRP3 | + | -10 | 2367 | 1 | | | 2 | | | | | |
| 2 | 82535054 | + | DHFRP3 | + | -7 | 2318 | 1 | | | | | | 2 | | |
| 2 | 108944368 | - | SULT1C2P1 | + | -1 | 2919 | 1 | | | | | | | 13 | |
| 2 | 118066845 | - | DDX18 | + | -6 | 2277 | 1 | | | | | | | | 4 |
| 2 | 127685890 | + | LOC101929926 | + | -1 | 2893 | 1 | 2 | | | | | | | |
| 3 | 9409844 | - | THUMPD3 | + | -8 | 2301 | 1 | | 2 | | | | | | |
| 3 | 39276481 | + | CX3CR1 | - | -3 | 2320 | 1 | | | | 2 | | | | |
| 3 | 59488570 | + | FHIT | - | -1 | 2919 | 1 | 2 | | | | | | | |
| 3 | 60531466 | + | FHIT | - | 3 | 2909 | 1 | | 2 | | | | | | |
| 3 | 98452053 | + | ST3GAL6 | + | -4 | 2320 | 1 | | 2 | | | | | | |
| 3 | 102905729 | + | MIR548AB | - | -5 | 2321 | 1 | | | | | | 2 | | |
| 3 | 102905746 | + | MIR548AB | - | -8 | 2308 | 1 | | 2 | | | | | | |
| 3 | 127146691 | - | LINC02016 | - | -13 | 2406 | 1 | | | | | | 2 | | |
| 3 | 141137159 | + | ZBTB38 | + | -13 | 2315 | 1 | 10 | | | | 6 | 10 | | |
| 3 | 144493929 | + | C3orf58 | + | 0 | 2913 | 1 | | | | 3 | | | | |
| 3 | 180142369 | + | LINC02053 | + | -3 | 2945 | 1 | | | | | 2 | | | |
| 4 | 6341261 | - | PPP2R2C | - | -11 | 2335 | 1 | | 2 | | | | | | |
| 4 | 16665581 | - | LDB2 | - | -10 | 2283 | 1 | | | | 2 | | | | |
| 4 | 91960654 | - | CCSER1 | + | 12 | 2909 | 1 | | 2 | | | | | | |
| 4 | 144322959 | + | GAB1 | + | 0 | 2915 | 1 | | | | 2 | | | | |
| 4 | 146192817 | + | OTUD4 | - | -7 | 2443 | 1 | | | | 2 | | | | |
| 4 | 163008230 | + | FSTL5 | - | 4 | 2908 | 1 | | 2 | | | | | | |
| 4 | 182561840 | + | LINC00290 | - | -2 | 2298 | 1 | | | | 2 | | | | |
| 5 | 531513 | + | MIR4456 | - | -10 | 2453 | 1 | | | | 2 | | | | |
| 5 | 531612 | + | MIR4456 | - | -1 | 2313 | 1 | | | 2 | | | | | |
| 5 | 1038588 | - | NKD2 | + | -12 | 2328 | 1 | | | | | 2 | 3 | | |
| 5 | 35857136 | + | IL7R | + | -1 | 2383 | 1 | 38 | | | | 230 | 42 | | |
| 5 | 103909276 | - | RAB9BP1 | + | 1 | 2909 | 1 | | | 2 | | | | | |
| 5 | 109050641 | - | MAN2A1 | + | -1 | 2923 | 1 | | 2 | | | | | | |
| 5 | 113472235 | + | KCNN2 | + | 1 | 2909 | 1 | 2 | | | | | | | |
| 5 | 174393316 | + | LINC01951 | - | -1 | 2312 | 1 | | | 2 | | | | | |
| 5 | 178864994 | + | ADAMTS2 | - | -10 | 2457 | 1 | | 2 | | | | | | |

| | | | | | | | | | | | | | | | |
|---|---|---|---|---|---|---|---|---|---|---|---|---|---|---|---|
| 5 | 180237145 | - | MGAT1 | - | -7 | 2322 | 1 | 2 | | | | | | | |
| 6 | 74230885 | - | EEF1A1 | - | -13 | 2329 | 1 | | | | | 6 | | | |
| 6 | 74230950 | - | EEF1A1 | - | -2 | 2383 | 1 | | | | | 187 | 25 | | |
| 6 | 86042776 | - | LOC101928820 | - | 4 | 2909 | 1 | | | | 3 | | | | |
| 7 | 21592931 | + | DNAH11 | + | -2 | 2320 | 1 | | | | 2 | | | | |
| 7 | 37569594 | + | ELMO1 | - | -16 | 2340 | 1 | 4 | | | | 6 | 7 | | |
| 7 | 41967580 | + | GLI3 | - | -7 | 2310 | 1 | 2 | | | | | | | |
| 7 | 54981095 | + | EGFR | + | -1 | 2909 | 1 | | 2 | | | | | | |
| 7 | 68787219 | - | LOC100507468 | - | -1 | 2945 | 2 | | | | 4 | | | | |
| 7 | 137087811 | - | DGKI | - | -7 | 2443 | 1 | | 2 | | | | | | |
| 7 | 142499043 | - | PRSS3P2 | + | 3 | 2913 | 2 | | | | 2 | | | | |
| 8 | 24735650 | + | NEFM | + | -11 | 2836 | 2 | | | | | 2 | | | |
| 8 | 104108859 | + | ATP6V1C1 | + | 0 | 2304 | 1 | | | | | | 3 | | |
| 8 | 111954749 | - | LINC01608 | - | 0 | 2299 | 1 | | | 4 | | | | | |
| 8 | 124970617 | + | FER1L6 | + | -2 | 2319 | 1 | | | | | | 2 | | |
| 8 | 143316756 | - | TSNARE1 | - | -6 | 2322 | 1 | | 2 | | | | | | |
| 9 | 15517752 | + | PSIP1 | - | -1 | 2918 | 2 | | 2 | | | | | | |
| 9 | 24241091 | + | IZUMO3 | - | -10 | 2305 | 1 | 2 | | | | | | | |
| 9 | 97136757 | + | MFSD14B | + | -3 | 2318 | 1 | | 2 | | | | | | |
| 9 | 140012353 | - | DPP7 | - | -1 | 2886 | 1 | 2 | | | | | | | |
| 10 | 7677856 | + | ITIH5 | - | -10 | 2314 | 1 | 3 | | | | | | | |
| 10 | 73513417 | - | VSIR | - | -4 | 2305 | 1 | | | 2 | | | | | |
| 10 | 73513422 | - | VSIR | - | -1 | 2300 | 1 | | 2 | | | | | | |
| 10 | 126312033 | - | FAM53B | - | -10 | 2339 | 1 | | 2 | | | | | | |
| 11 | 2016269 | - | HOTS | + | -9 | 2321 | 1 | | | | | | 2 | | |
| 11 | 8113994 | + | TUB | + | 0 | 2909 | 1 | | | 2 | | | | | |
| 11 | 45628130 | - | CHST1 | - | -19 | 2338 | 1 | | | | 2 | | | | |
| 11 | 70717804 | - | SHANK2 | - | 8 | 2909 | 1 | | 6 | | | | | | |
| 11 | 72136216 | + | CLPB | - | -1 | 2910 | 1 | | | | | 9 | | | |
| 11 | 76512155 | - | TSKU | + | -5 | 2310 | 1 | | 2 | | | | | | |
| 11 | 76804291 | - | CAPN5 | + | -11 | 2333 | 1 | 2 | | | | | | | |
| 11 | 93248241 | - | SMCO4 | - | 8 | 2940 | 1 | | | | | | | | 2 |
| 11 | 106583926 | + | GUCY1A2 | - | -8 | 2296 | 1 | | | | 2 | | | | |
| 11 | 126427530 | - | KIRREL3 | - | -8 | 2276 | 1 | | | | | | 2 | | |
| 12 | 7540051 | + | CD163L1 | - | -6 | 2282 | 1 | 2 | | | | | | | |
| 12 | 50909869 | - | DIP2B | + | -5 | 2305 | 1 | | | | | 2 | | | |
| 12 | 66994805 | - | GRIP1 | - | -8 | 2308 | 1 | 3 | | | | | 2 | | |
| 12 | 66994834 | - | GRIP1 | - | -4 | 2277 | 1 | | | 2 | | | | | |
| 12 | 117814318 | + | NOS1 | - | -11 | 2327 | 1 | | | | | | 2 | | |
| 12 | 133055408 | + | FBRSL1 | + | -14 | 2330 | 1 | | 2 | | | | | | |
| 13 | 58282758 | - | PCDH17 | + | -7 | 2312 | 1 | 13 | | | | 2 | 7 | | |
| 13 | 58282809 | - | PCDH17 | + | -11 | 2374 | 1 | | | | | 2 | | | |
| 13 | 74020397 | - | LINC00392 | + | -6 | 2332 | 1 | | | | | | 2 | | |

| | | | | | | | | | | | | | | | |
|---|---|---|---|---|---|---|---|---|---|---|---|---|---|---|---|
| 14 | 23016469 | + | DAD1 | - | 5 | 2376 | 1 | | | | | | 7 | | |
| 14 | 35874560 | - | NFKBIA | - | -11 | 2328 | 1 | | 2 | | | | | | |
| 14 | 58166114 | + | SLC35F4 | - | -3 | 2917 | 1 | | | | | 2 | | | |
| 15 | 38969247 | + | C15orf53 | + | 2 | 2934 | 1 | | | 2 | | | | | |
| 15 | 61849059 | + | VPS13C | - | -5 | 2276 | 1 | | | | | 2 | | | |
| 15 | 68892594 | + | CORO2B | + | -18 | 2285 | 1 | | | | 2 | | | | |
| 15 | 101538839 | + | LRRK1 | + | -4 | 2320 | 1 | | 3 | | | | | | |
| 15 | 101538843 | + | LRRK1 | + | -4 | 2320 | 1 | | | | | | 3 | | |
| 16 | 20320088 | + | GP2 | - | -9 | 2304 | 1 | | | 2 | | | | | |
| 16 | 87475882 | - | ZCCHC14 | - | -12 | 2325 | 1 | | 2 | | | | | | |
| 16 | 87680328 | + | JPH3 | + | -12 | 2328 | 1 | | | | | | 4 | | |
| 16 | 88725273 | + | MVD | - | -9 | 2452 | 1 | | | | | | 2 | | |
| 16 | 88834811 | + | PIEZO1 | - | -12 | 2328 | 1 | | | | 2 | | | | |
| 17 | 5312929 | - | NUP88 | - | -2 | 2302 | 1 | | | | 5 | | | | |
| 17 | 6681539 | - | FBXO39 | + | -7 | 2297 | 1 | | 2 | | | | | | |
| 17 | 21787830 | - | FAM27E5 | + | 12 | 2909 | 1 | | 2 | | | | | | |
| 17 | 36860489 | + | MLLT6 | + | -7 | 2323 | 1 | | | | | | 2 | | |
| 17 | 40433369 | + | STAT5B | - | -5 | 2323 | 1 | 2 | | | | | 2 | | |
| 17 | 57763859 | - | CLTC | + | 7 | 2919 | 1 | | 3 | | | | | | |
| 17 | 72885258 | + | FADS6 | - | -6 | 2449 | 1 | | 2 | | | | | | |
| 18 | 5096328 | + | AKAIN1 | - | 0 | 2883 | 1 | | 2 | | | | | | |
| 18 | 20036927 | - | CTAGE1 | - | 16 | 2938 | 1 | | 2 | | | | | | |
| 18 | 22690653 | - | ZNF521 | - | -3 | 2941 | 1 | | | | | 3 | | | |
| 18 | 22703920 | + | ZNF521 | - | -1 | 2929 | 2 | | | | | 7 | | | |
| 18 | 64807084 | + | MIR5011 | + | 1 | 2899 | 1 | | | 2 | | | | | |
| 19 | 412347 | - | C2CD4C | - | -11 | 2454 | 1 | | | | | | | | 2 |
| 19 | 13879366 | + | MRI1 | + | -8 | 2328 | 1 | | | | | | 4 | | |
| 19 | 13879583 | + | MRI1 | + | -10 | 2334 | 1 | | 2 | | | | | | |
| 19 | 29987082 | + | LOC284395 | - | -8 | 2332 | 1 | | | | 2 | | | | |
| 19 | 37765032 | + | LOC284412 | - | -11 | 2449 | 1 | | | 2 | | | | | |
| 19 | 49665405 | - | TRPM4 | + | -4 | 2298 | 1 | | | 3 | | | | | |
| 19 | 52771813 | - | ZNF766 | + | -3 | 2913 | 2 | | | | 2 | | | | |
| 20 | 2901461 | + | PTPRA | + | -1 | 2888 | 1 | 2 | | | | | | | |
| 20 | 26189072 | + | MIR663AHG | - | 1 | 2909 | 1 | | | | | | 3 | | |
| 20 | 33451666 | + | GGT7 | - | -18 | 2333 | 1 | 2 | | | | | | | |
| 20 | 36361690 | - | CTNNBL1 | + | 9 | 2909 | 1 | | 2 | | | | | | |
| 20 | 36361706 | + | CTNNBL1 | + | -1 | 2884 | 1 | | 3 | | | | | | |
| 20 | 42845486 | + | OSER1-AS1 | + | -6 | 2442 | 1 | | | 2 | | | | | |
| 20 | 47122175 | - | PREX1 | - | -2 | 2318 | 1 | | 2 | | | | | | |
| 20 | 54404096 | - | CBLN4 | - | -7 | 2328 | 1 | 2 | | | | | | | |
| 21 | 46572295 | + | ADARB1 | + | -9 | 2312 | 1 | | | | 2 | | | | |
| 22 | 26320985 | + | MYO18B | + | -1 | 2896 | 1 | | 2 | | | | | | |
| X | 48516388 | + | WAS | + | -3 | 2926 | 1 | 2 | | | | | | | |

| | | | | | | | | | | | | | | | |
|---|---|---|---|---|---|---|---|---|---|---|---|---|---|---|---|
| X | 48541559 | - | WAS | + | 22 | 2909 | 1 | | 5 | | | | | | |
| X | 48541821 | - | WAS | + | -8 | 2908 | 1 | | | | | | 2 | | |
| X | 48541835 | - | WAS | + | 0 | 2909 | 1 | | 4 | | | | | | |
| X | 48541838 | - | WAS | + | 0 | 2923 | 1 | 3 | | | | | | | |
| X | 48541922 | - | WAS | + | -4 | 2914 | 1 | | | | | | | | 2 |
| X | 48542247 | + | WAS | + | 3 | 2909 | 1 | | | | | | | 3 | |
| X | 48542247 | - | WAS | + | -1 | 2919 | 1 | | | | | | 5 | | |
| X | 48542248 | + | WAS | + | -1 | 2910 | 1 | | | | 2 | | | | |
| X | 48542642 | + | WAS | + | -1 | 2915 | 2 | | | | | 5 | | | |
| X | 48542647 | + | WAS | + | 9 | 2909 | 1 | | | | | | 51 | | |
| X | 48542654 | + | WAS | + | -4 | 2920 | 2 | | | | 6 | | | | |
| X | 48542657 | + | WAS | + | 16 | 2925 | 1 | 2 | | | | | | | |
| X | 48542659 | + | WAS | + | -2 | 2925 | 1 | | 3 | | | | | | |
| X | 48543692 | - | WAS | + | -4 | 2915 | 2 | | | | 6 | | | | |
| X | 48547310 | + | WAS | + | -3 | 2889 | 1 | 2 | | | | | | | |
| X | 48552110 | + | SUV39H1 | + | 7 | 2889 | 1 | 2 | | | | | | | |

**Table S4. Most targeted genes by AAV integration.**

| ALL IS N= 164 | | | |
|---|---|---|---|
| chr | GeneID | Targeting | % |
| chrX | WAS | 16 | 9.76 |
| chr1 | SMPDL3B | 7 | 4.27 |
| chr1 | KDM4A | 3 | 1.83 |
| chr2 | DHFRP3 | 3 | 1.83 |
| chr10 | VSIR | 2 | 1.22 |
| chr12 | GRIP1 | 2 | 1.22 |
| chr13 | PCDH17 | 2 | 1.22 |
| chr15 | LRRK1 | 2 | 1.22 |
| chr18 | ZNF521 | 2 | 1.22 |
| chr19 | MRI1 | 2 | 1.22 |
| chr20 | CTNNBL1 | 2 | 1.22 |
| chr3 | FHIT | 2 | 1.22 |
| chr3 | MIR548AB | 2 | 1.22 |
| chr5 | MIR4456 | 2 | 1.22 |
| chr6 | EEF1A1 | 2 | 1.22 |

| CD34 N=115 | | | |
|---|---|---|---|
| chr | GeneID | Targeting | % |
| chrX | WAS | 10 | 8.7 |
| chr1 | SMPDL3B | 3 | 2.61 |
| chr1 | KDM4A | 3 | 2.61 |
| chr10 | VSIR | 2 | 1.74 |
| chr12 | GRIP1 | 2 | 1.74 |
| chr2 | DHFRP3 | 2 | 1.74 |
| chr20 | CTNNBL1 | 2 | 1.74 |
| chr3 | FHIT | 2 | 1.74 |
| chr5 | MIR4456 | 2 | 1.74 |

| BM=57 | | | |
|---|---|---|---|
| chr | GeneID | Targeting | % |
| chrX | WAS | 6 | 10.71 |
| chr1 | SMPDL3B | 4 | 7.14 |
| chr13 | PCDH17 | 2 | 3.57 |
| chr18 | ZNF521 | 2 | 3.57 |
| chr6 | EEF1A1 | 2 | 3.57 |

### Table S5. AAV integration sites abundance in SU samples in vitro (relative to Figure 6G)

| SU-1 CD34 | | | SU-2CD34 | | | SU-3 CD34 | | | SU-4 CD34 | | |
|---|---|---|---|---|---|---|---|---|---|---|---|
| GeneID | % | Cell count | GeneID | % | Cell count | GeneID | % | Cell count | GeneID | % | Cell count |
| IL7R | 30.4 | 38 | SHANK2 | 6.7 | 6 | ATP6V1C2 | 30.9 | 17 | WAS | 8.2 | 6 |
| PCDH17 | 10.4 | 13 | WAS | 5.6 | 5 | LINC01608 | 7.3 | 4 | WAS | 8.2 | 6 |
| ZBTB38 | 8 | 10 | WAS | 4.5 | 4 | SMPDL3B | 5.4 | 3 | NUP88 | 6.8 | 5 |
| SMPDL3B | 4 | 5 | LRRK1 | 3.4 | 3 | TRPM4 | 5.4 | 3 | LOC100507468 | 5.5 | 4 |
| ELMO1 | 3.2 | 4 | CLTC | 3.4 | 3 | OBSCN | 3.6 | 2 | C3orf58 | 4.1 | 3 |
| KIF17 | 2.4 | 3 | CTNNBL1 | 3.4 | 3 | VSIR | 3.6 | 2 | LOC101928820 | 4.1 | 3 |
| KDM4A | 2.4 | 3 | WAS | 3.4 | 3 | TUB | 3.6 | 2 | KDM4A | 2.7 | 2 |
| ITIH5 | 2.4 | 3 | RABGGTB | 2.2 | 2 | GRIP1 | 3.6 | 2 | LINC00466 | 2.7 | 2 |
| GRIP1 | 2.4 | 3 | SEC22B | 2.2 | 2 | C15orf53 | 3.6 | 2 | PKN2-AS1 | 2.7 | 2 |
| WAS | 2.4 | 3 | VSIR | 2.2 | 2 | GP2 | 3.6 | 2 | EVI5 | 2.7 | 2 |

**Table S6. AAV integration sites abundance tracking in vivo for SU-1 (relative to Figure 6H)**

| SU-1 CD34 | | | SU-1 17BM | | | SU-1 21BM | | |
|---|---|---|---|---|---|---|---|---|
| GeneID | % | Cell count | GeneID | % | Cell count | GeneID | % | Cell count |
| IL7R | 30.4 | 38 | IL7R | 42.6 | 230 | WAS | 24.3 | 51 |
| PCDH17 | 10.4 | 13 | EEF1A1 | 34.6 | 187 | IL7R | 20 | 42 |
| ZBTB38 | 8 | 10 | NCF2 | 8.2 | 44 | EEF1A1 | 11.9 | 25 |
| SMPDL3B | 4 | 5 | HFM1 | 2.4 | 13 | ZBTB38 | 4.8 | 10 |
| ELMO1 | 3.2 | 4 | CLPB | 1.7 | 9 | PCDH17 | 3.3 | 7 |
| KIF17 | 2.4 | 3 | ZNF521 | 1.3 | 7 | DAD1 | 3.3 | 7 |
| KDM4A | 2.4 | 3 | ZBTB38 | 1.1 | 6 | ELMO1 | 3.3 | 7 |
| ITIH5 | 2.4 | 3 | EEF1A1 | 1.1 | 6 | WAS | 2.4 | 5 |
| GRIP1 | 2.4 | 3 | ELMO1 | 1.1 | 6 | JPH3 | 1.9 | 4 |
| WAS | 2.4 | 3 | WAS | 0.9 | 5 | MRI1 | 1.9 | 4 |

### Table S7. AAV integration sites abundance tracking in vivo for SU-3 (relative to Figure 6I)

| **SU-3 CD34** | | | **SU-3 27BM** | | | **SU-3 28BM** | | |
|---|---|---|---|---|---|---|---|---|
| GeneID | % | Cell count | GeneID | % | Cell count | GeneID | % | Cell count |
| ATP6V1C2 | 30.91 | 17 | SULT1C2P1 | 65 | 13 | DDX18 | 26.67 | 4 |
| LINC01608 | 7.27 | 4 | ATP6V1C2 | 20 | 4 | ECHDC2 | 20 | 3 |
| SMPDL3B | 5.45 | 3 | WAS | 15 | 3 | SMCO4 | 13.33 | 2 |
| TRPM4 | 5.45 | 3 | | | | C2CD4C | 13.33 | 2 |
| OBSCN | 3.64 | 2 | | | | ATP6V1C2 | 13.33 | 2 |
| VSIR | 3.64 | 2 | | | | WAS | 13.33 | 2 |
| TUB | 3.64 | 2 | | | | | | |
| GRIP1 | 3.64 | 2 | | | | | | |
| C15orf53 | 3.64 | 2 | | | | | | |
| GP2 | 3.64 | 2 | | | | | | |
